# The conserved aphid saliva chemosensory protein effector Mp10 targets plant AMSH deubiquitinases at cellular membranes to suppress pattern-triggered immunity

**DOI:** 10.1101/2024.11.15.622802

**Authors:** Matteo Gravino, Sam T. Mugford, Nathan Kreuter, Joshua Joyce, Christine Wilson, Adi Kliot, James Canham, Thomas C. Mathers, Claire Drurey, Abbas Maqbool, Carlo Martins, Gerhard Saalbach, Saskia A. Hogenhout

**Affiliations:** Crop Genetics Dept., John Innes Centre, Norwich Research Park, Norwich, NR4 7UH, United Kingdom; Biochemistry and Metabolism Dept., John Innes Centre, Norwich Research Park, Norwich, NR4 7UH, United Kingdom; Proteomics Technology Platform, John Innes Centre, Norwich Research Park, Norwich, NR4 7UH, United Kingdom

## Abstract

Chemosensory proteins (CSPs) are a conserved family present in insects and other arthropods, recognized for their critical roles in both intra- and interspecies communication. However, the functional mechanisms of these proteins remain largely unexplored. In our previous research, we identified a CSP in aphid saliva, Mp10, from the peach-potato aphid *Myzus persicae*, which functions as an effector protein modulating host plant immunity. Mp10 suppresses pattern recognition receptor (PRR)-triggered immunity (PTI), the first layer of plant defence, while also inducing effector-triggered immunity (ETI). In this study, we elucidate the molecular mechanisms by which Mp10 suppresses PTI. Our findings reveal that Mp10 interacts with AMSH deubiquitinase enzymes in plants, as shown by yeast two-hybrid, co-immunoprecipitation (co-IP), and FRET-FLIM assays, with these interactions predominantly localized to intracellular membranes. Mp10 was found to modulate the dynamics of membrane-bound PRR receptor kinases in plant cells. Co-IP and mass spectrometry analyses demonstrated that Mp10 and AMSH2 associate with a range of PRR kinases, PRR-associated kinases, and proteins involved in the intracellular trafficking of membrane proteins. Mp10 reduces the accumulation of these kinases at the cell surface by promoting their internalization to internal membranes, thereby dampening PTI. Supporting this, a dominant-negative catalytically inactive variant of AMSH2 also inhibits PTI. Interestingly, Mp10 orthologues from other sap-feeding hemipteran insects exhibit similar immune-suppressive activities, and our findings show that their interaction with plant AMSH proteins is conserved, indicating this immune-suppression mechanism is evolutionarily ancient.

## Introduction

Chemosensory proteins (CSPs) form a conserved family found in insects and other arthropods (1), playing significant roles in communication within species, such as in social insects, and across species (2). For example, a fly CSP is involved in direct recognition of fungal virulence factors, triggering behavioural defences that benefit fly survival against infection (3). Blood-feeding Diptera, such as mosquitoes, inject anti-hemostatic and anti-inflammatory D7 proteins, which contain an OBP domain structurally related to CSPs (4), to aid in blood feeding (5). Likewise, aphids inject a CSP, known as Mp10, into plant cells at the onset of feeding (6). While Mp10 triggers plant defences (7, 8), aphids likely deliver this protein into plant cells to modulate specific processes for their benefit, as Mp10 also suppresses the plant initial defence response (7). However, mechanisms involved in CSP immune suppression is unknown.

Aphids (Hemiptera, Sternorrhyncha, Aphidoidea) are major agricultural pests. Effective host perception is crucial for their survival and feeding success. Aphids feed primarily on phloem sap using specialized piercing-sucking mouthparts (stylets) that facilitate the injection of saliva and uptake of cell contents (9). Upon landing on a plant, prior to reaching the phloem, aphids probe different cell types and tissues during an initial exploratory phase. This early interaction allows them to assess whether establishing a long-term feeding relationship with the host vasculature system or abandon it in search of a more suitable plant. During this early probing phase, aphids may also introduce plant viruses (10). Most aphid species are highly specialized, rejecting most plants they encounter, while a few species, like the green peach aphid *Myzus persicae* (Sulzer), have an unusually broad host range among insect herbivores (11). How aphids make these feeding decisions remains unclear, but it likely involves an intricate balance between plant resistance factors that restrict feeding and aphid virulence factors that modulate plant defences.

Plant immune systems have two well understood layers. Firstly, PAMP-triggered immunity (PTI) occurs when plant cell surface pattern recognition receptors (PRRs) recognize conserved pathogen- or herbivore-associated molecular patterns (PAMPs). Adapted pathogens can suppress PTI via the delivery of effector proteins into plant hosts. In a second line of defence, plants possess intracellular nucleotide-binding leucine-rich repeat (NLR) immune receptors that recognise the pathogen/pest effectors or their activities leading to effector-triggered immunity (ETI), often a stronger immune response than PTI, triggering cell death or other defence mechanisms that limit pathogen spread or insect feeding (12). The robustness of the plant immune system is maintained through crosstalk between the PTI and ETI branches of immunity (13), ensuring effective protection against a wide array of threats.

Given that Mp10 can both suppress and activate plant defences (7, 8, 14, 15), it may play a critical role in aphid host selection and acceptance. We previously found that Mp10 is delivered into the cytoplasm of plants cells, including mesophyll cells, during the initial stages of aphid feeding (6). Mp10 specifically binds to the acrostyle (16), a structure at the tip of aphid stylets that bind viruses (17). From this position, Mp10 can be directly released into the plant cell cytoplasm when the aphid stylets probe the plant cells (6). Mp10 suppresses the plant initial defence response (6, 7). However, Mp10 also triggers plant immune responses, causing chlorosis resembling cell death-like features in leaves, and leading to a significant reduction in aphid reproduction (7, 8, 14). Since several major effectors of plant pathogens both suppress and activate plant defences (18–20), we hypothesized that Mp10 plays a role in modulating key plant processes involved in plant-aphid interactions.

In this study we uncovered how Mp10 suppresses PTI. We found that Mp10 interacts with plant AMSH deubiquitinase enzymes at intracellular membranes to modulate PRR receptor kinase homeostasis. AMSH proteins are conserved across eukaryotes and play important roles in the regulation of cell-surface receptors in animals (21) and PRR receptors in plants (22). Interestingly AMSH proteins are also involved in the regulation of the intracellular NLR receptors involved in immune signalling in animals (23) and ETI in plants (24). Mp10 causes the mis-localisation and reduces accumulation of PRR proteins, rendering plant cells insensitive to elicitors and suppressing PTI. Similar effects are seen with Mp10 orthologues from other sap-feeding hemipteran pests that also suppress PTI and interact with plant AMSH proteins, suggesting a 250-million-year history of interaction with plant AMSH.

## Results

### Mp10 orthologues from sap-feeding hemipteran insects suppress PTI

It was previously shown that Mp10 suppresses the plant PTI response to the bacterial PAMP flg22, a peptide derived from bacterial flagellin (7), and also to aphid-derived elicitors (25). Moreover, orthologues from other sap-feeding hemipterans also possess the same PTI-suppressing activity (25). CSP proteins were identified in *Acyrthosiphon pisum* (pea aphid), *Diuraphis noxia* (Russian wheat aphid), *Aphis gossypii* (cotton/melon aphid), *Brevicoryne brassicae* (cabbage aphid), *Diaphorina citri* (Asian citrus psyllid), and *Bemisia tabaci* (tobacco whitefly), from the Sternorrhyncha suborder, as well as *Nilaparvata lugens* (rice brown planthopper), *Macrosteles quadrilineatus* (Aster leafhopper), *Circulifer tenellus* (beet leafhopper), *Dalbulus maidis* (corn leafhopper), from the Auchenorrhyncha suborder (Fig 1A, adapted from our previously published preprint manuscript (25)). When transiently produced in *Nicotiana benthamiana* leaves as N-terminal eGFP-fusion proteins, MpCSP4 (Mp10), AgCSP4, ApCSP4, BtCSP4, CtCSP4 and DmCSP4 suppressed the extracellular ROS and Ca^2+^ burst responses to flg22 (Fig 1B, 1C and 1D, adapted from our previously published preprint manuscript (25)). These data suggest that the origin of the plant immune-suppressive activity possessed by the CSP4 clade evolved before the divergence of the Sternorrhyncha and Auchenorrhyncha sub-orders of sap-feeding hemipteran herbivores.

**Fig 1.**
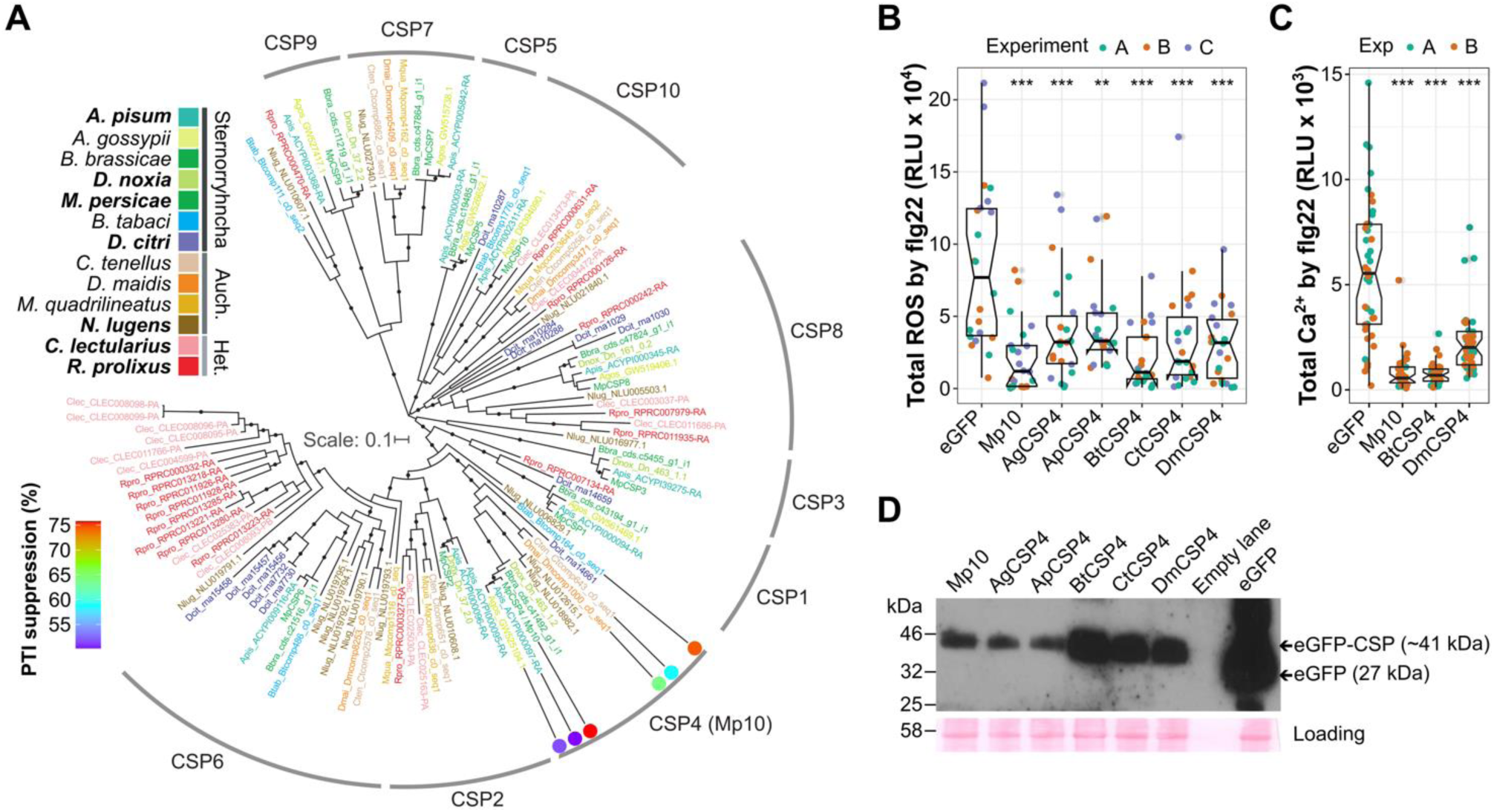
Mp10 (CSP4) orthologs are conserved among sap-feeding hemipterans and suppress plant immune responses. Adapted from (25). (**A**) Mp10 forms a distinct clade with CSP orthologs from sap-feeding hemipterans. The maximum likelihood phylogenetic tree, rooted at the mid-point, shows species color-coded by lineage, with bold text for those with available whole genome sequences. Circles indicate SH-like support values >0.8. Colored dots represent CSPs tested for PTI suppression, with colors indicating the level of suppression (legend, lower left). Auch., Auchenorrhyncha; Het., Heteroptera. (**B**) Mp10 orthologs suppress ROS bursts in *N. benthamiana* WT leaf disks producing eGFP or eGFP-fusion CSPs upon flg22 exposure (100 nM). Y-axis: total ROS burst (RLU) over 60 minutes; X-axis: eGFP-alone (control) and eGFP-CSP fusion proteins from various hemipteran species. (**C**) Mp10 orthologs suppress Ca^2+^ bursts, which were measured using a bioluminescent reporter assay in *N. benthamiana* transgenic plants expressing eGFP-aequorin and eGFP or eGFP-fusion CSPs upon flg22 exposure. Y-axis: total Ca^2+^ burst (RLU) over 30 minutes; X-axis: eGFP-alone (control) and eGFP-CSP fusion proteins from select species. In (B-C), box plots show median, interquartile range (IQR), and whiskers denote 1.5*IQR. Data points are from individual leaf disks across three independent experiments. Asterisks indicate significant differences from the control (ANOVA, Tukey HSD, **p < 0.01; ***p < 0.001). (**D**) Protein expression verification for ROS assays in (B). Western blot detects EGFP in leaf disks expressing eGFP or eGFP-fusion CSPs. MW markers are indicated on the left; expected bands are marked by arrows. Protein loading is shown using Ponceau S staining.

### Mp10 orthologues target the conserved JAMM domain of AMSH deubiquitinating enzymes

To identify molecular targets of Mp10 in the host plant, we conducted a yeast two-hybrid (Y2H) screen with Mp10 against an *Arabidopsis thaliana* (hereafter Arabidopsis) cDNA library. Plasmids were isolated from 6 colonies that showed growth on selective media, these were re-transformed together with the pLexA-Mp10 or the empty pLexA bait vector. Three out of 6 showed growth in the presence of the Mp10 bait vector but not the empty vector. All three clones contained an insert including the full-length coding sequence of ASSOCIATED MOLECULE WITH THE SH3 DOMAIN OF STAM 2 (AMSH2, At1g10600).

AMSH2 is a JAMM-domain (JAB1/MPN/MOV34 metalloenzyme domain, IPR000555) containing putative de-ubiquitinating enzyme (DUB) activity that belongs to a small family with three members in Arabidopsis (26). AMSH1 and AMSH3 associate with components of the endosomal sorting complex required for transport (ESCRT) and play important roles in the regulation of membrane- and membrane-protein trafficking (22, 26). Interestingly, the ESCRT complexes are involved in the trafficking and regulation of PRR-kinases such as the FLAGELLIN-SENSING 2 (FLS2) receptor that recognizes flg22 (27), suggesting a mechanism by which Mp10 might suppress the sensitivity of these receptors to elicitors. To date, the function of AMSH2 in plants has remained largely unclear.

AMSH proteins are conserved across higher plants, with most plant species having one orthologue of each AMSH1, 2, and 3 per haploid genome (26). We cloned the coding sequences of AMSH1, AMSH2, and AMSH3 from Arabidopsis Col-0 into yeast-2-hybrid plasmids and found evidence that Mp10 also interacts with AMSH1 and AMSH3 (Fig 2A). The interaction of Mp10 with AMSH2 was detected under higher stringency conditions and promoted greater growth of yeast than with AMSH1 or AMSH3, consistent with a stronger interaction with AMSH2 (Fig 2A). Some autoactivation was observed with AMSH3 – i.e., the AMSH3-DNA-binding domain (BD) fusion was sufficient to promote growth on selective media in the presence of DNA activating domain (AD)- alone – but this growth was always substantially weaker than in the presence of Mp10-AD, indicating a likely interaction (Fig 2A). Because of the conserved plant immune-suppressive activity of Mp10 with orthologues from other sap-feeders, and the similar conservation of the AMSH proteins in plant genomes, we hypothesised that the interaction of Mp10 with AMSH proteins might be similarly conserved. First, we found that Mp10 interacts with AMSH2 orthologues from other dicot plant species, including *Brassica napus* (rapeseed), *Citrus sinensis* (sweet orange), *Pisum sativum* (pea), *Beta vulgaris* (beet), and *N. benthamiana* (Fig 2A). Moreover, we found that the Mp10 orthologues from other sap-feeding insects, including whiteflies (*B. tabaci*) and leafhoppers (*C. tenellus*, *M. quadrilineatus*), also interacted with AMSH2 from Arabidopsis (Fig 2B). The conservation of interaction between Mp10 orthologues and Arabidopsis AMSH2 is consistent with the ancient origin of the Mp10 immune-suppressive activity (Fig 1). Y2H assays together with the phylogenetic analyses suggest that the Mp10/CSP4 clade acquired the binding specificity to the conserved JAMM domain of AMSH before the divergence of plant-feeding hemipterans, indicating that the interaction is ancient and conserved among sap-feeding hemipterans.

**Fig 2.**
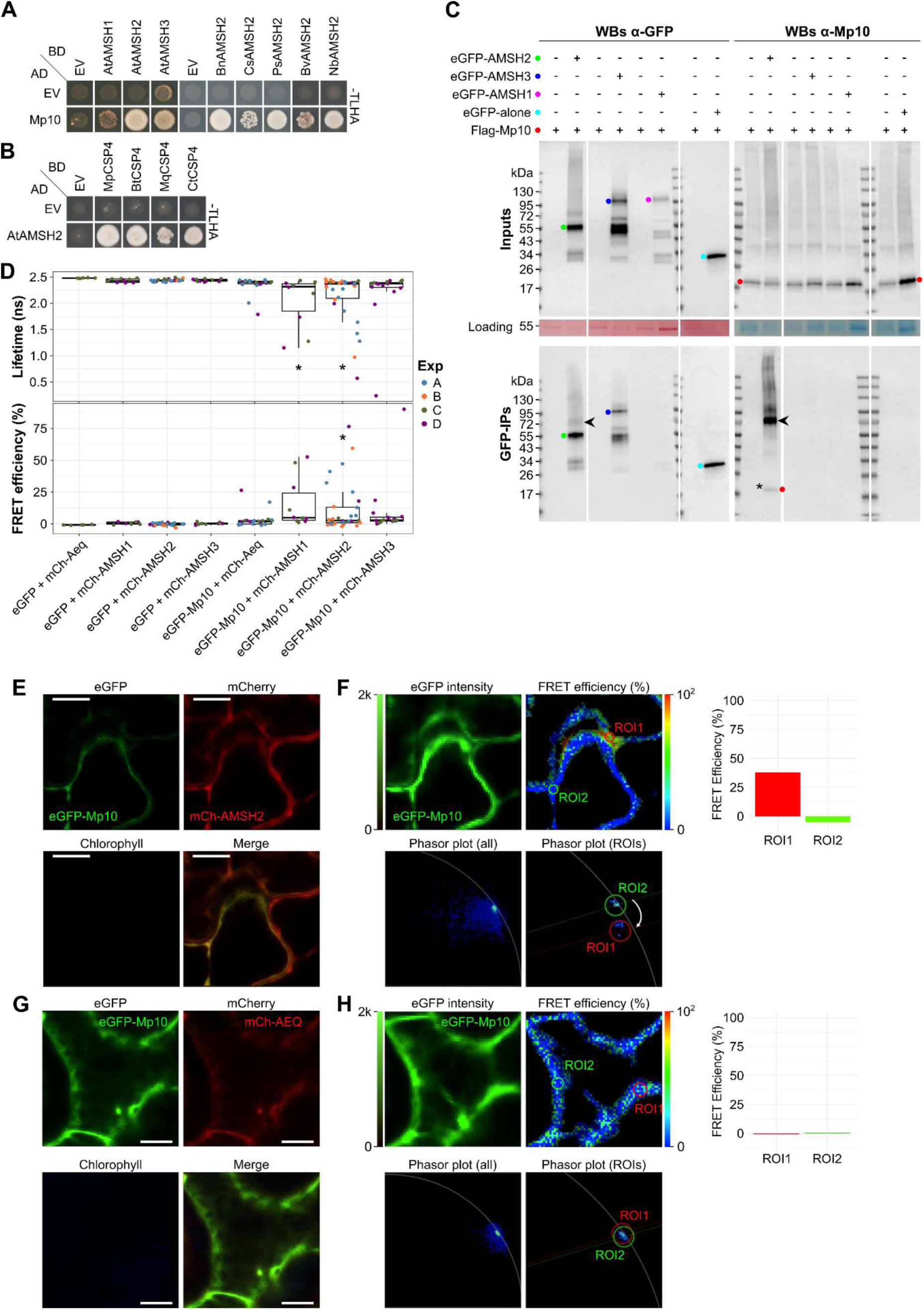
Mp10 interacts with AMSH deubiquitinases, at distinct plant subcellular loci. (**A-B**) Yeast two-hybrid (Y2H) assays show interactions between MpCSP4/Mp10 and AMSH1, AMSH2, and AMSH3 from Arabidopsis and AMSH2 from other plant species (**A**), and CSP4 orthologs from various insects with AMSH2 from Arabidopsis (**B**). EV: empty vector; AD: GAL4-activation domain; BD: GAL4-DNA binding domain; -TLHA: quadruple dropout medium. (**C**) AMSH2 pulls down Mp10 from *N. benthamiana* membrane fractions. Co-immunoprecipitation (co-IP) were conducted from *N. benthamiana* leaves transiently expressing Flag-Mp10 with(out) eGFP-alone (control) or eGFP tagged Arabidopsis AMSH1, AMSH2, AMSH3, followed by western blot analyses using antibodies to eGFP and Mp10 (see S2A Fig for Flag detection). Protein sizes indicated by molecular weight markers (kDa). Expected bands marked with colored dots; asterisk denotes Flag-Mp10 band pulled down with AMSH2; arrowheads denote a putative modified form of Mp10, and/or an oligomeric state of Mp10, or a higher-order complex containing the Mp10 protein that is apparently resistant to denaturation by SDS-PAGE. Loading visualized with Ponceau S or Coomassie Blue staining. (**D**) FLIM analysis in *N. benthamiana* leaves co-expressing eGFP-Mp10 or eGFP-alone (as control) with mCherry-tagged AMSHs or aequorin (AEQ, as control). The fluorescence lifetime (top) and derived FRET efficiency (bottom) of eGFP donor fluorophores alone or fused to Mp10 (y-axis) in presence of mCherry acceptor fluorophores fused to AMSHs or AEQ (x-axis). Each datapoint is the average for a whole image from a different cell (n = 5 to 31) across 4 biological replicate experiments. Asterisk (*) indicates significant difference from eGFP-alone + mCherry-AMSH2 control (p-adj <0.05, one-way ANOVA with Tukey HSD). Boxplots display median, 25th/75th percentiles, and data range across four independent experiments. (**E-H**) Observation of high FRET efficiency between mCherry-AMSH2 and eGFP-Mp10 in specific subcellular cytoplasmic regions of interest (ROIs). At left, confocal images of eGFP-Mp10 with mCherry-AMSH2 (E) and eGFP-Mp10 with mCherry-AEQ (control) (G). At right, FRET analysis of eGFP-Mp10 with mCherry-AMSH2 (F) and eGFP-Mp10 with mCherry-AEQ (control) (H) with panels showing eGFP-Mp10 intensity (upper left), FRET efficiency heat maps (upper right), and phasor plots for full images (lower left) and selected ROIs (lower right). Graphs to the right show the extracted FRET efficiency values (y-axis) for the selected ROIs (x-axis).

### Mp10 interacts AMSH proteins at specific subcellular locations within plant cells

To assess if Mp10 and AMSH interact *in planta*, we performed a GFP co-immunoprecipitation (co-IP) assay from *N. benthamiana* leaves co-infiltrated with constructs producing eGFP-tagged Arabidopsis AMSH2 and Flag-tagged Mp10 protein. GFP co-IP from a total soluble protein extract did not pull down Mp10 with AMSH2 at the expected molecular weight (S1 Fig). However, the Flag antibody detected higher molecular weight proteins (about 80 kDa, <175 kDa, and >175 kDa, compared to the band of ∼18 kDa expected for Flag-Mp10) that co-immunoprecipitated with eGFP-AMSH2 only in the presence of Flag-Mp10, and not with eGFP-alone (S1 Fig). Although AMSH proteins do not directly associate with membranes, they are recruited by other proteins into membrane-associated complexes that regulate membrane protein turnover (26, 28). Therefore, we examined whether Mp10 interacts with Arabidopsis AMSHs in a membrane-associated manner. Membrane fraction enrichment by ultracentrifugation followed by GFP co-IP (29) co-immunoprecipitated Flag-Mp10 at the expected molecular weight with eGFP-AMSH2, but not the control Flag-RFP (Figs 2C, S2A and S2B). Co-immunoprecipitation of Flag-Mp10 was not detected with eGFP-AMSH3, eGFP-alone or GFP beads alone (Figs 2C and S2A). The co-IP of Flag-Mp10 and eGFP-AMSH1 was inconclusive, because eGFP-AMSH1 expression was low and not detectable upon GFP co-IP on western blots (Fig 2C). These results suggest that Mp10 and AMSH2 interact *in planta*, most likely at cellular membranes.

In addition to a protein of ∼18 kDa that corresponds to Flag-Mp10 in the eGFP-AMSH2 pull down from membrane-enriched fractions, the Mp10 and Flag antibodies reacted with proteins of ∼80 kDa, and a ladder of higher molecular weight proteins (Figs 2C and S2A), similar to that observed in the co-IP from total soluble protein extract (S1 Fig). This band could represent a modified form of Mp10, an oligomeric state of Mp10, or a higher-order complex containing the Mp10 protein that is apparently resistant to denaturation by SDS-PAGE, as has been observed for other membrane-associated complexes (30), and reduction of disulphide bonds by DTT. A band at ∼80 kDa is detected also with the anti-GFP antibody in the Co-IP of eGFP-AMSH2 plus Flag-Mp10, but not eGFP-AMSH2 plus Flag-RFP (Figs 2C and S2B), indicating that eGFP-AMSH2 might be part of the same SDS-resistant complex. These data show that Mp10 and AMSH2 may be part of a larger complex that is resistant to denaturation, a feature often associated with membrane-associated complexes (30).

To further investigate the complex formation between Mp10 and AMSH2 and its subcellular localization in plant cells, we conducted *in vivo* fluorescence lifetime imaging (FLIM)-Förster resonance energy transfer (FRET) assays. Since not all the Arabidopsis AMSHs showed high and detectable levels of expression when transiently produced in *N. benthamiana* (see Fig 2C), we used *N. benthamiana* AMSHs for these experiments. First, eGFP-Mp10 and eGFP alone (control) as donor fluorophores were co-produced with mCherry-AMSH2 or mCherry-aequorin (control) as acceptor fluorophores in *N. benthamiana* leaves. Then, the donor fluorophores were excited at 488 nm, and their lifetimes were measured using a Leica Stellaris microscope in time-correlated single photon counting mode. Figure 2D shows the modelled fluorescence lifetimes and FRET efficiency averaged across whole images including cytoplasm of cells expressing both fluorescent protein fusions; each data point comes from a separate cell (n = 5 to 31) across 4 independent experiments. We observed that the lifetime of eGFP-Mp10 was significantly reduced only in the presence of mCherry-AMSH2, and not with mCherry-aequorin (Fig 2D), while the lifetime of eGFP alone was similar in all samples (Fig 2D). FRET efficiency values were modelled from the FLIM data and mapped onto the confocal images. eGFP-Mp10 and mCherry-AMSH2 showed co-localization throughout the cytosol (Fig 2E). We observed high FRET efficiency at isolated subcellular locations in the presence of eGFP-Mp10 together with mCherry-AMSH2. In Fig 2F ROI1 indicates an area of high FRET efficiency compared to ROI2 in the same image, indicating that Mp10 and AMSH2 interact with each other at specific cellular localizations, consistent with localizations of complexes at membranes. No such areas of high FRET were detected in cells co-expressing eGFP-Mp10 and mCherry-aequorin, despite largely co-localizing in the cytosol (Fig 2G and 2H). Phasor plot analysis provides an alternative measure of changes in fluorescence lifetime caused by FRET (31), a shift in a clockwise direction on a phasor plot is indicative of protein-protein interaction. We observed that the regions of high FRET efficiency in the presence of eGFP-Mp10 together with mCherry-AMSH2 showed a clockwise shift relative to an area of low FRET efficiency (ROI1 *vs* ROI2 in Fig 2F). No such shifts in the phasor plots of eGFP-Mp10 in the presence of mCherry-aequorin were seen (Fig 2 H). These data support the modelled changes in fluorescence lifetime and FRET efficiency, indicating protein-protein interaction between eGFP-Mp10 and mCherry-AMSH2 at certain locations.

Additionally, FLIM-FRET detected an interaction of eGFP-Mp10 with mCherry-AMSH1 but not with mCherry-AMSH3 (Figs 2D, S3A and S3B). mCherry-AMSH1 showed a distinct subcellular localization, concentrating in discrete spots, approximately one per cell and 10 µm in size, rather than being distributed throughout the cytosol like mCherry-AMSH2. In these loci, eGFP-Mp10 partially co-localized with mCherry-AMSH1 with high FRET efficiency and a corresponding clockwise shift in the phasor plot (S3A and S3B Fig). Some areas of high FRET efficiency were observed at different locations compared to the observed location of mCherry-AMSH1 (for example ROI3 in S3B Fig), this is likely due to the movement of the mCherry-AMSH1 during the FLIM data gathering which took several seconds. The subcellular localization pattern of mCherry-AMSH3 was mainly similar to that of mCherry-AMSH2 (Figs 2E and S3C) and occasionally showing discrete loci similar to mCherry-AMSH1 in some cells (S3A and S3E Fig), but was not associated with a significant change in fluorescence lifetime of eGFP-Mp10 resulting in increased FRET efficiency in either case (S3D and S3F Fig). Therefore, Mp10 does not seem to interact with AMSH3 within plant cells, in agreement with the co-IP results. Overall, these results suggest that Mp10 most likely physically interacts with AMSH2 and AMSH1 in isolated subcellular loci within plant cells.

### AMSH2 and Mp10 form complexes *in vitro*

We also evaluated the direct interaction by isolating complexes between Mp10 and AMSH2. First, we performed an *in vivo* glutathione S-transferase (GST) co-immunoprecipitation (Co-IP) assay in bacteria. For this assay, GST fusion proteins of AMSH2 from Arabidopsis and GFP-binding protein (GBP) were co-expressed with or without Mp10 and immunoprecipitated from *Escherichia coli* protein extracts. Mp10 was co-immunoprecipitated with GST-AMSH2, but not with GST-GBP (Fig 3A). In addition, Mp10 also interacted with AMSH1 and AMSH3 (Fig 3A). Therefore, Mp10 interacts with all three AMSH proteins in the *E. coli* extract.

**Fig 3.**
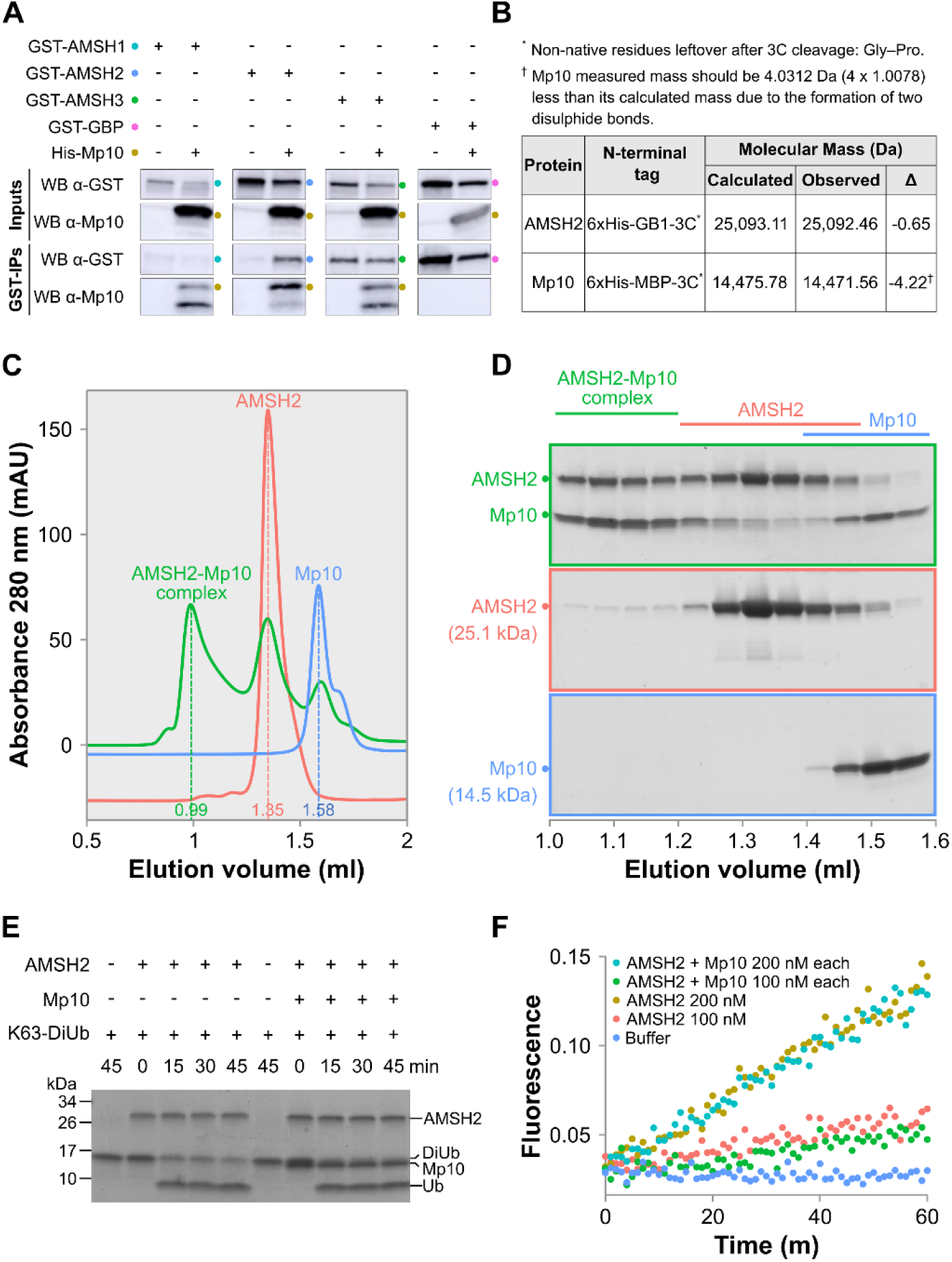
Mp10 forms a complex with AMSHs *in vitro*. (**A**) GST pull-down assays showing interactions of GST-tagged AMSH1, AMSH2, AMSH3 with His-tagged Mp10, and not GBP. Proteins were detected on western blots using anti-GST and anti-Mp10 antibodies. Expected protein bands are marked with colored dots. (**B**) Intact mass spectrometry of purified Mp10 and AMSH2 proteins (see S4A Fig) confirms a Mp10 observed mass ∼4 Da lower than the calculated mass, consistent with two disulfide bonds. (**C-D**) Gel filtration assays showing Mp10 directly interacts with Arabidopsis AMSH2. (**C**) Chromatographic profiles display UV absorbance at 280 nm (mAU) versus elution volume (ml), with peak elution volumes marked. (**D**) Coomassie-stained SDS-PAGE gels of fractions collected from fractions in (C). Expected protein bands are indicated with colored dots. (**E-F**) *In vitro* deubiquitination (DUB) assays using K63-linked di-ubiquitin substrates, unlabeled (E) or labeled with TAMRA (F), in the presence or absence of AMSH2 alone or AMSH2 with Mp10. In (E), Coomassie-stained gels show reactions at specified time points, with MW markers on the left and expected bands on the right. Additional DUB assays are shown in S4B and S4C Fig. In (F), quantitative DUB activity measurements, recorded as fluorescence changes every minute over 60 minutes, are shown for various conditions including protein buffer (negative control), USP2 (positive control), AMSH2, and AMSH2 with Mp10.

To verify Mp10-AMSH2 complexes, we purified these proteins for *in vitro* gel filtration assays. The purification process involved immobilized metal ion affinity chromatography (IMAC) followed by high-resolution preparative size exclusion chromatography (SEC). After removing the tag using 3C protease, AMSH2 and Mp10 underwent further purification with reverse IMAC and SEC. The expected size of AMSH2 and Mp10 (25,093 Da and 14,476 Da, respectively) was confirmed by Coomassie staining and intact mass spectrometry analysis (Figs 3B and S4A). The experimentally observed mass of Mp10 was approximately 4 Da lower than the theoretical mass, aligning with the predicted formation of two disulfide bonds in Mp10 (25). Purified Mp10 and AMSH2 were then mixed in equimolar amounts and analysed by analytical size-exclusion chromatography (SEC), with each protein alone serving as a control. As expected, analytical SEC of Mp10 and AMSH2 individually showed distinct peaks corresponding to their separate elution fractions, confirmed by Coomassie staining (Fig 3C and 3D). However, when the proteins were combined, in addition to the individual peaks, a third, more prominent peak appeared at a reduced retention volume (Fig 3C), indicating the formation of a complex. Coomassie staining confirmed that the new peak contained both Mp10 and AMSH2 within the same fractions (Fig 3D). Altogether, these data demonstrate that Mp10 and AMSH2 directly interact and form a complex *in vitro*.

### Mp10 does not modify AMSH2 DUB activity towards di-ubiquitin substrates *in vitro*

AMSH1 and AMSH3 have been shown to have DUB activity toward K63-linked ubiquitin chains (26, 32), which is associated with initiation of protein endocytosis (33). However, that of AMSH2 has not yet been investigated. Here, we found that AMSH2 is an active enzyme that possesses DUB activity toward K63-linked di-ubiquitin, either unlabelled or labelled with a fluorescent reporter (TAMRA) and a highly efficient quenching dye pair (Fig 3E and 3F). However, in our experiments the presence of Mp10 did not affect AMSH2 DUB activity, either at the same molar concentration (Fig 3E and 3F) or with two or four times Mp10 molar concentration compared to AMSH2 (S4B Fig). Because we found that there is a proportion of unbound AMSH2 in the AMSH2-Mp10 complex sample (see Fig 3C and 3D), which may buffer the Mp10-AMSH2 complex DUB activity, we also tested the activity of the purified fraction containing only the AMSH2-Mp10 complex (Fig 3C and 3D) compared to AMSH2-alone (Fig 3C and 3D). Also in this condition, we did not find differences in AMSH2 DUB activity between samples (S4C Fig). These results indicate that in these *in vitro* conditions, Mp10 does not affect the AMSH2 DUB activity towards K63-linked di-ubiquitin substrates.

### DUB enzymatic activity of AMSH2 is required for PTI

AMSH deubiquitinase (DUB) activity is crucial for proper plant development and the effector-induced degradation of ubiquitinated membrane receptor kinases (22, 32). We discovered that Mp10 suppresses PTI and interacts with AMSH2 in plants; however, the role of AMSH2 DUB activity in PTI signalling remains unclear. To investigate this, we transiently expressed mCherry-tagged wild-type AMSH2, an enzymatically inactive AMSH2 variant with an active site AXA mutation (H127A, H129A) (32), and mCherry alone (as a control), along with eGFP-Mp10 or eGFP alone, in *N. benthamiana*. We then assessed ROS bursts in response to the bacterial elicitor flg22 (7). Our assays showed a significant reduction in ROS production in treatments that included eGFP-Mp10, mCherry-AMSH2(AXA) and both eGFP-Mp10 and mCherry-AMSH2(AXA) (Fig 4A and 4B). Additive effects when both were present were not observed (Fig 4A), indicating they operate in the same pathway. Moreover, mCherry-AMSH2 did not suppress ROS production, whereas mCherry-AMSH2 + eGFP-Mp10 did (Fig 4A and 4B). These results show that both AMSH2(AXA) and Mp10 suppress PTI, consistent with the Mp10-mediated suppression of immune-responses operating via the interaction with the AMSH2 protein.

**Fig 4.**
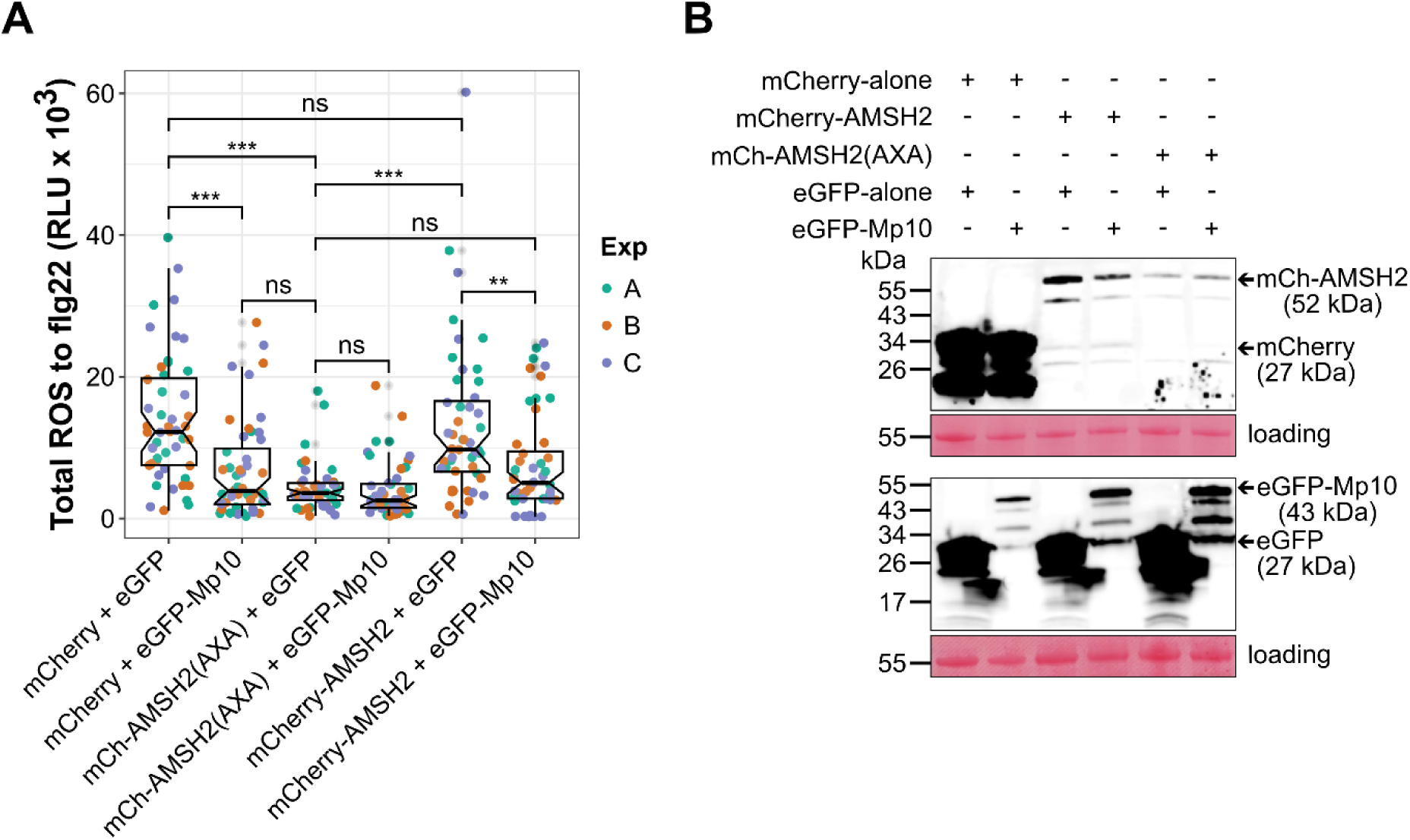
Mp10 and the enzymatically inactive AMSH2 mutant (AMSH2-AXA) suppress PTI. (**A**) Luminol-based ROS burst assay in *N. benthamiana* leaf disks co-expressing mCherry alone, mCherry-AMSH2, or AMSH2(AXA) with either eGFP alone (control) or eGFP-Mp10, following exposure to 100 nM flg22. Y-axis: total ROS burst (RLU) over 60 minutes; X-axis: co-expressed constructs. Box-plots display median, interquartile range (IQR), and whiskers denote 1.5*IQR. Data points represent individual leaf discs, with colors indicating independent replicates. Asterisks denote significant differences (ANOVA, Tukey HSD; *p < 0.05; **p < 0.01; ***p < 0.001; ^ns^not significant). (**B**) Protein expression verification for ROS assays. Western blots detect mCherry and eGFP in leaf disks expressing mCherry variants and eGFP constructs. MW markers are shown on the left; expected bands are marked by arrows. Protein loading was visualized using Ponceau S staining.

### Stable Mp10-overexpressing lines and *amsh2* homozygous null mutants exhibit lethality

To investigate the potential effects of AMSH2 and Mp10 on *M. persicae* interactions with plants, we aimed to generate *amsh2* null mutants and stable Mp10 overexpression lines. A previously reported Arabidopsis AMSH2 T-DNA insertion mutant in the 5’ UTR of the AMSH2 gene (CSHL_ET4018) was found to exhibit no significant reduction in AMSH2 transcript levels (26). We identified another T-DNA insertion mutant (SALKseq_055175.1) annotated as having an insertion in the AMSH2 coding region, but the mutant allele was combined with a wild-type allele of AMSH2 in a heterozygous line, and our attempts to isolate homozygous lines were unsuccessful. Then, we attempted to generate a knock-out mutant via CRISPR. Heterozygous transgenic plants were generated with a 668-base pair deletion spanning a region from the 1^st^ to the 3^rd^ exons of AMSH. 192 T2 offspring from 16 T1 plants were screened for the presence of the deletion mutant, but only heterozygous or WT offspring were recovered at a ratio of 2:1 indicating that the homozygous AMSH2 knock-out is lethal. We generated eGFP-AMSH2 overexpressing lines. These showed high level of eGFP-AMSH2 expression, without an obvious effect on aphid fecundity (S5A and S5B Fig), in agreement with our finding that transient overexpression of AMSH2 in plants does not affect PTI (see Fig 4A).

We also attempted to generate Mp10 overexpressing lines. An N-terminal eGFP fusion of Mp10 was cloned into the binary vector pB7WGF2 under the control of 35S promoter and subsequently transformed into Arabidopsis Col-0 WT plants by floral dip. On average, we obtained 0.667 (± 0.21 SEM) Basta-resistant T1 transformants per pot of dipped plants (n = 6). However, none of the surviving transgenics showed detectable levels of Mp10 protein expression, indicating that stable overexpression of Mp10 is lethal.

### Immune receptor proteins and proteins involved in intracellular transport are co-immunoprecipitated with AMSH2 and Mp10

Mp10 suppresses the PRR receptor kinase mediated responses to a variety of elicitor molecules (7, 25), and AMSH proteins regulate the homeostasis of membrane-bound receptor kinases (21, 22, 34). We hypothesised that the membrane-associated interaction of Mp10 with AMSH2 might involve PRR receptor kinase proteins and affect their homeostasis. To test this, we performed reciprocal co-immunoprecipitation experiments of eGFP-AMSH2 and Flag-Mp10 from membrane enriched fractions from *N. benthamiana* tissue producing one or both proteins and analysed the precipitated proteins by mass spectrometry (see Fig 2C).

Through the analysis of co-IP fractions from three independent experiments, we identified significantly enriched proteins with a fold change in abundance greater than 4 and adjusted p-value lower than 0.05. This revealed 17 proteins that co-immunoprecipitated exclusively with eGFP-AMSH2 in the presence of Mp10—including Mp10 itself—and 53 proteins in its absence, compared to eGFP-alone controls (S6A, S6B and S6C Fig) (35). Of these, 6 proteins were common to both conditions (S6A Fig) (35).

In a reciprocal analysis, we performed Mp10 pull downs and mass spectrometry on co-IP samples of Flag-Mp10/Flag-alone with eGFP-AMSH2, and Flag-Mp10/Flag-alone with eGFP-alone. We identified 46 proteins exclusively co-immunoprecipitated with Flag-Mp10 in the absence of eGFP-AMSH2, and 7 proteins in its presence, including eGFP-AMSH2, compared to Flag-alone controls (S6A, S6D and S6E Fig) (35). Among these, 1 protein was common to both conditions (S6A Fig) (35). Notably, we did not detect the endogenous *N. benthamiana* AMSH proteins in the pull-down with Mp10, despite having shown we can detect the interaction of NbAMSH2 and NbAMSH1 with Mp10 by FLIM-FRET (see Figs 2D, 2E, 2F, S3A and S3B). Possibly, the endogenous expression levels of AMSHs are too low to reliably detect them by mass spectrometry in this experimental set up.

Notably, in the eGFP-AMSH2 pulldown in the presence of Flag-Mp10 there was a significant decrease - approximately threefold - in the number of unique proteins co-immunoprecipitated compared to eGFP-AMSH2 alone (S6A Fig). Similarly, in the Mp10 pull down in the presence of eGFP-AMSH2 there was a significant decrease – approximately sevenfold - in the number of unique proteins co-immunoprecipitated compared to Flag-Mp10 alone (S6A Fig). These data suggest that Mp10 may inhibit AMSH2 interaction with certain proteins, redirecting it to engage with other complexes instead.

In the Flag-Mp10 plus eGFP-AMSH2 pull-down assays, we identified Nbe.v1.s00110g27840 annotated as a RLK similar to AT1G1490 (fold change = 19.251 relatively to Flag-alone plus eGFP-AMSH2, adjusted p-value = 0.000855), whereas the Flag-Mp10 without eGFP-AMSH2 identified Nbe.v1.s00070g28750 annotated as lectin-type receptor kinase S5 similar to AT5G06740 (fold change = 6.256 relative to Flag-alone plus eGFP-alone, adjusted p-value = 0.017156) (S1 Table) (35). In addition, we found 5 further receptor kinase proteins that pull down with eGFP-AMSH2 either in the presence or absence of Mp10, albeit all when applying a significance threshold without a false-discovery rate control - given the abundance of this protein class within the list, it is likely that most are true discoveries rather than false positives (S2 Table) (35). In an initial analysis of the data using an earlier version of Proteome Discoverer software—where proteomics outcomes can be notably affected by the choice of analytical tools, especially for low-abundance proteins (36)— Nbe.v1.s00060g27420, an orthologue of FERONIA (FER) from Arabidopsis (AT3G51550), was identified as co-purifying with AMSH2 in an Mp10-dependent manner. In another co-IP experiment with a single experimental replicate, FER was also pulled-down with eGFP-AMSH2 in the presence of Mp10 (5.6-fold enriched) but not in the absence of Mp10 (0.4-fold enriched) (35). FER has known regulatory functions affecting the activity of a multitude of other receptor-mediated processes, including those involving FLS2 (36). As Mp10 reduces the sensitivity of plant cells to elicitors acting via receptor kinase proteins, and the AMSH proteins are involved in the degradation of receptors, we hypothesised that the receptor kinases we tentatively identified as pulling down with AMSH or Mp10 might be targeted for degradation, and thus be present only at low abundance rendering the detection of the interaction with Mp10 and AMSH more challenging.

In addition to receptor kinases, we identified proteins involved in the trafficking of membranes and membrane proteins around the cell, including a Syntaxin (Nbe.v1.s00180g20930) that co-IP with Mp10 in presence of AMSH2 (fold change = 5.766 and adjusted p-value = 0.00023) (S1 Table) (35). In addition, we found a further Syntaxin and a coatomer subunit that pull down with eGFP-AMSH2 in the presence of Mp10, as well as a vesicle-associated membrane protein (VAMP) that pulls down with eGFP-AMSH2 either in the presence or absence of Mp10, and a number of Rab GTPAses - albeit all when applying a significance threshold without a false-discovery rate control (S1 and S2 Tables) (35). All proteins that regulate the fusion of membranes and the trafficking of membrane proteins between cellular compartments (37–40). Similarly to FER, in an early analysis of the data using a previous version of the Proteome discoverer software, we found that Nbe.v1.s00050g03900 an orthologue of Arabidopsis vacuolar protein sorting 2.1 (VPS2.1, AT2G06530) co-IPs with Mp10, although in this analysis the fold-change is below the 4-fold threshold (S1 Table) (35). The VPS2.1 protein is a component of the ESCRT-III complex known to interact with AMSH1 and AMSH3 (22, 26), and is involved in the degradation of ubiquitinated membrane proteins (26).

### Mp10 affects the abundance and localisation of cell-surface receptor-like kinases

FER, one of the receptor-like kinase that was tentatively detected as co-immunoprecipitated with AMSH2 in presence of Mp10, is a malectin-like receptor kinase that facilitates the ligand-induced complex formation of the immune receptor kinase FLS2 with their co-receptor BAK1 to initiate immune signalling (36, 41). Given that we previously found that Mp10 suppresses flg22-mediated induction of PTI via FLS2 (7), we reasoned that Mp10 may affect the homeostasis of FER and FLS2. To test this hypothesis, we co-expressed Arabidopsis FER as a C-terminal RFP fusion alongside Flag-Mp10 or Flag-alone in *N. benthamiana* leaves, and then assessed the abundance of the proteins by western blot (WB) analysis. In absence of Mp10, a band corresponding to FER-RFP expected size (∼128 kDa) was detected on the blot (Fig 5A). However, in presence of Mp10 the abundance of FER-RFP was drastically reduced (Fig 5A), suggesting that Mp10 mediates the destabilization of FER. Similarly, Arabidopsis FLS2 as a C-terminus eGFP-tagged fusion destabilized in the presence of Mp10 in *N. benthamiana* leaves (Fig 5B). In contrast, the presence of Mp10 did not have any effect on the accumulation of the Arabidopsis CERK1 chitin receptor fused with 6xHA or 1xHA tags at its C-terminus (S7 Fig), in agreement with Mp10 not suppressing chitin-induced PTI (7).

**Fig. 5.**
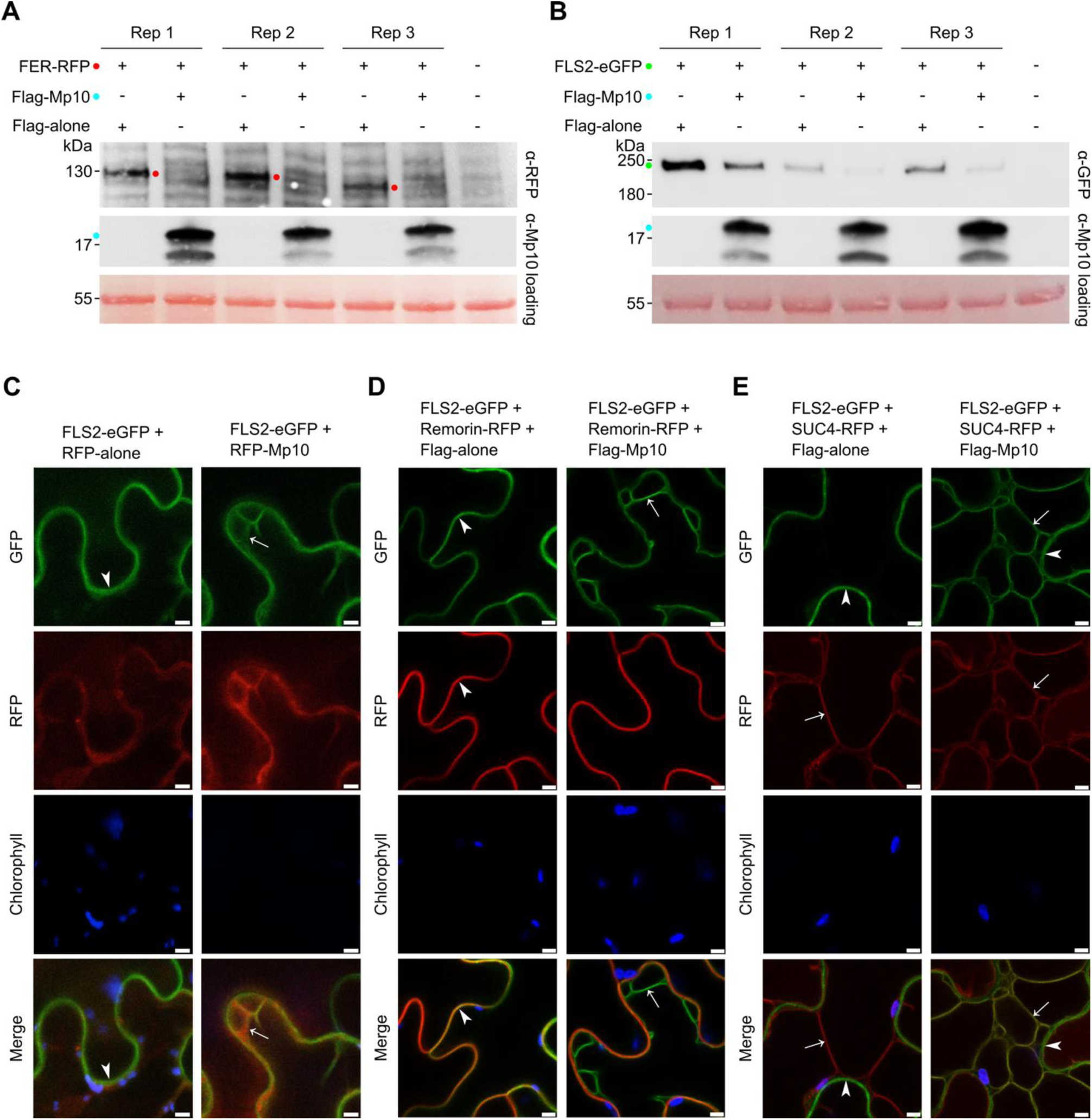
Mp10 affects the abundance and localisation of cell-surface receptor-like kinases. **(A)** Western blot analyses showing Arabidopsis FERONIA (FER)-RFP protein stabilization in presence/absence of Mp10 using antibodies to RFP (top panel) or Mp10 (middle panel). (**B**) Western blot analyses of Arabidopsis FLS2-eGFP protein stabilization in presence/absence of Mp10 using antibodies to eGFP (top panel) or Mp10 (middle panel). In (**A-B**), expected bands marked with colored dots. Loading visualized with Ponceau S staining. (**C**) Confocal microscopy analyses showing tomato FLS2 subcellular localization in presence of RFP-Mp10 or RFP-alone (control). (**D-E**) Confocal microscopy analyses showing tomato FLS2 subcellular localization in presence of Flag-Mp10 or Flag-alone (control). Remorin-RFP labels the plasma membrane, as indicated by arrowheads. StSUC4-RFP labels the tonoplast membrane, as indicated by arrows.

The degradation pathway of FLS2 involves its endocytosis via the endosome pathway, progressing through multi-vesicular bodies to the lumen of the vacuole (27, 42). We performed confocal microscopy experiments to examine the subcellular localization of FLS2-eGFP in the presence of RFP-Mp10. When co-expressed with RFP-alone, the FLS2-eGFP signal was exclusively observed at the outer edge of the cytoplasm (Fig 5C), as defined by the cytoplasmic RFP signal from RFP-alone (43), consistent with the plasma membrane localization of FLS2 (44). The RFP signal from RFP-Mp10 is localized in the cytoplasm, as reported previously (14). However, when co-expressed with RFP-Mp10, the overall signal of FLS2-eGFP was weaker. In cells with stronger signal, we observed an unusual localisation pattern, with FLS2-eGFP present both at the outer edge and the inner edge of the cytoplasm (Fig 5C). Additionally, the FLS2-eGFP signal was observed along the inner edge of chloroplasts and surrounding what appear to be transvacuolar cytoplasmic strands (Fig 5C) - i.e. tubes of cytoplasm that traverse the vacuole – an observation not seen in the absence of Mp10. The PM-localized marker protein Remorin (45) and the tonoplast-localized marker protein StSUC4 (46) were used to further identify the membranes. We found that in absence of Flag-Mp10, FLS2-eGFP co-localized with Remorin but not with StSUC4 (Fig 5D and 5E). However, in presence of Flag-Mp10, FLS2-eGFP co-localize with both Remorin and StSUC4 (Fig 5D and 5E). These data suggest that, in absence of Mp10, FLS2-eGFP is exclusively localised in the plasma membrane. In contrast, in presence of Mp10, FLS2-eGFP mis-localizes to also locate at vacuolar membranes. These results indicate that Mp10 can affect the homeostasis and localization of RLKs involved in PTI signalling.

## Discussion

In this study, we characterized the function of the CSP Mp10, present in aphid saliva (6). We show that Mp10 suppresses the plant initial defence response, known as PTI, aligning with observations that aphids deposit this CSP directly into the cytoplasm of mesophyll cells (6), which aphids typically probe during the early phase of feeding (7). We demonstrate that Mp10 binds to conserved plant AMSH deubiquitinases. The interaction of Mp10 and AMSH2 is direct as shown by yeast two-hybrid, co-immunoprecipitation (co-IP), and FRET-FLIM assays, and predominantly localized to intracellular membranes within plant cells, consistent with known roles of AMSH proteins as part of membrane-associated protein complexes (47). Moreover, Mp10 and AMSH2 form a complex *in vitro*. In the presence of AMSH2, Mp10 precipitates with membrane-associated receptor proteins, including RLKs/RLPs and proteins involved in the trafficking of membrane proteins. We found that Mp10 destabilizes key cell-surface immune receptors FLS2 and FER that both form a complex with the co-receptor BRASSINOSTEROID INSENSITIVE 1-ASSOCIATED KINASE 1 (BAK1) (36), previously shown to contribute to plant immunity to aphids (48), to mediate flg22-induced ROS and Ca^2+^ bursts of plants (49). In agreement with this, we observed that Mp10 promotes localization of FLS2 to intracellular membranes, providing an explanation of how Mp10 suppresses flg22-induced ROS and Ca^2+^ bursts of plants. Altogether, these results suggest that Mp10 recruits AMSH2 to cellular membranes leading to re-localization and destabilization of cell-surface immune receptors and suppression of PTI (Fig. 6).

**Fig. 6.**
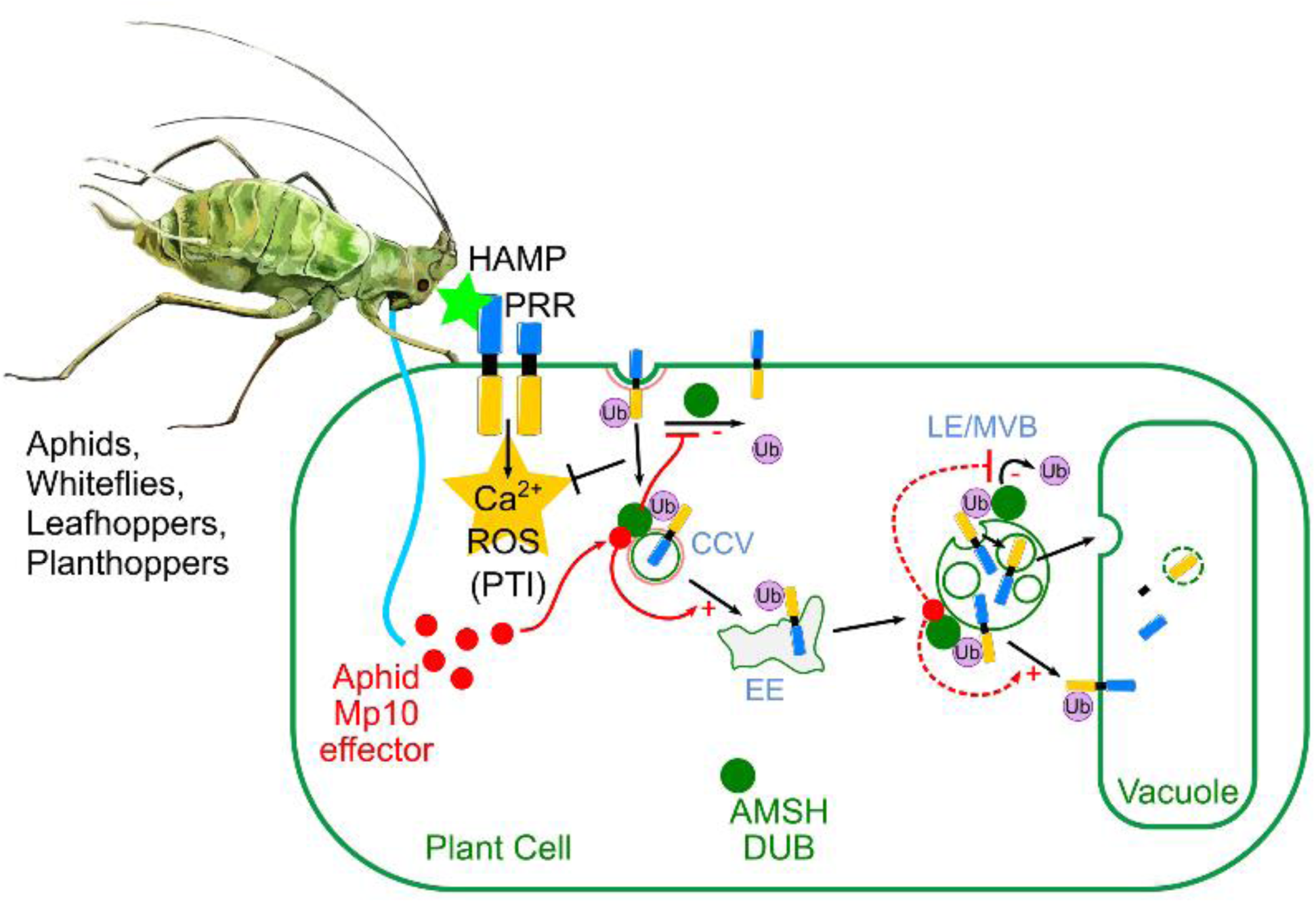
Model: Mp10 acts as local anesthetic to suppress pattern-triggered immunity (PTI). The aphid Mp10 effector is delivered into plant cells during aphid feeding. Mp10 binds to AMSH (DUB) enzymes at cellular membranes. We hypothesize that, when the enzyme is in complex with Mp10, the DUB activity of AMSH toward ubiquitinated membrane-bound pattern recognition receptors (PRRs) is blocked either at the plasma membrane or early endosome to promote their internalization and degradation-explaining the reduced abundance of receptor kinases in the presence of Mp10- and at the late endosome/multi-vesicular body to prevent internalization of PRRs into the intraluminal vesicles -explaining the observation of FLS2-GFP on the tonoplast membrane in the presence of Mp10. As consequence, in presence of Mp10 the PRRs are targeted for degradation in the endocytic pathway and/or re-localized into intracellular membranes, thereby suppressing PTI. This immune-suppressive ability of Mp10 and its interaction with plant AMSH are conserved across orthologs from diverse hemipteran insects, suggesting an evolutionary adaptation over 250 million years ago. HAMP-herbivore-associated molecular pattern; PRR-pattern recognition receptor; PTI-pattern-triggered immunity; Mp10-aphid salivary protein; AMSH-deubiquitinase (DUB) enzymes; CCV-clathrin-coated vesicle; EE-early endosome; LE/MVB-late endosome/multivesicular body. The model was generated using the following icons (https://bioicons.com/): Plant_cells icon by SwissBioPics https://www.swissbiopics.org/ is licensed under CC-BY 4.0 Unported https://creativecommons.org/licenses/by/4.0/; aphid icon by DBCLS https://togotv.dbcls.jp/en/pics.html is licensed under CC-BY 4.0 Unported https://creativecommons.org/licenses/by/4.0/; cell_group icon by JhonnyXC https://github.com/JRider16 is licensed under CC0 https://creativecommons.org/publicdomain/zero/1.0/; channel-membrane-red-2 icon by Servier https://smart.servier.com/ is licensed under CC-BY 3.0 Unported https://creativecommons.org/licenses/by/3.0/. All the material in 4.0 license version was modified.

Mp10 interacts most strongly with AMSH2 and, to a lesser extent, with AMSH1, while showing no interaction with AMSH3 in plant cells. Here, we show that AMSH2, a relatively understudied protein compared to AMSH1 and AMSH3, possesses DUB activity. Our findings indicate that a catalytically inactive AMSH2(AXA) mutant suppresses plant PTI responses, mimicking the PTI suppression phenotype of Mp10, despite Mp10 not suppressing AMSH DUB activity under *in vitro* conditions. Whereas overexpression of wild-type AMSH2 does not increase ROS production of plants in response to flg22 or improve resistance to aphids, Mp10 suppresses the ROS response in the presence of AMSH2. Additionally, AMSH2 is essential for plant viability, as homozygous AMSH2 knock-outs are lethal, and lethality also occurs in Mp10 overexpression lines. Given that Mp10 and AMSH2 directly interact, the most plausible explanation of these data is that Mp10 affects AMSH2 activity *in planta*.

AMSH proteins are known to deubiquitinate membrane-bound cargo proteins tagged for degradation on the late endosome membrane, a process associated with the internalisation of the cargo during the formation of the multi-vesicular body (MVB) (50). AMSH2 is highly conserved among plants, and AMSH1 and AMSH3 likely arose from a eudicot-specific gene duplication event, or possibly from the loss of AMSH1 in monocots (26). AMSH3 is critical for vacuole formation, autophagosome accumulation, and vacuolar protein transport in *Arabidopsis thaliana* (32), while AMSH1 plays a key role in autophagic degradation, aiding plant survival, senescence regulation, and pathogen defence (26). Both proteins contain MIT domains that bind to the ESCRT-III subunit VPS2.1, with AMSH3 also interacting with ESCRT-I proteins (51). While the MIT domain is missing in AMSH2, our work suggests that this plant protein has also an important role in the trafficking of membrane proteins.

In humans, AMSH proteins are proposed to function at two distinct stages of endosomal sorting of plasma membrane proteins (52). In the early stage, AMSH may counteract the activity of E3 ligases that tag plasma membrane proteins for degradation, thereby stabilizing these proteins. At a later stage, AMSH is thought to facilitate the degradation of cargo proteins by deubiquitinating them at MVB or late endosomes. In plants, different AMSH homologues might similarly regulate these stages. The disruption of both processes by Mp10 would be expected to lead to the FLS2 protein remaining on the external membrane of the MVB, and thus when the MVB fuses with the vacuole, FLS2 would be integrated into the tonoplast rather than entering the vacuole lumen for degradation, consistent with the mis-localisation of FLS2 at vacuolar membranes we observed. Conversely, disruption of the early endosomal processes would likely cause FLS2 to enter the vacuole lumen directly, leading to its degradation, which is consistent with the destabilization of FLS2 we observed (Fig. 6). Alternatively, Mp10 action may direct the DUB activity of AMSH2 against specific targets including RKs to mis-regulate their trafficking and homeostasis. Interestingly, Mp10 is not the only aphid effector implicated in membrane-trafficking processes in plants. *M. persicae* effector Mp1, was shown to target VPS52, a protein involved in vesicle trafficking (53).

The co-evolution of parasite virulence factors and their targets in host organisms is often a dynamic evolutionary arms race (54). Data reported herein revealed that Mp10 CSP effector protein is ancient and conserved across sap-feeding hemipteran insects, and similarly targets a conserved plant protein with an essential function, AMSH2. However, Mp10 not only suppresses PTI through AMSH2, making the plant more susceptible to aphid attack, but also induces ETI (8). A recent study found that a CSP from the rice-specialist planthopper *Nilaparvata lugens*, called NlCSP11, triggers ETI in *N. benthamiana* by directly interacting with the NLR protein RCSP, a member of the TIR-NBS-LRR clade that mediates ETI signalling through the immune regulator EDS1 (8). Mp10-induced chlorosis and stunting in *N. benthamiana* were also found to depend on RCSP and EDS1. However, *M. persicae* colonizes *N. benthamiana* successfully (55, 56). Moreover, *M. persicae* delivers the Mp10 protein into plant cells during feeding (6), but no chlorosis or ETI is observed in this context. We hypothesise that during aphid feeding, other factors contribute to the successful suppression of plant immunity, and that perhaps effector proteins exist in aphid saliva that suppress ETI act in concert with Mp10 to ensure the establishment of a feeding site. This aligns with the role of *M. persicae* effector Mp64, which targets E3 SUMO ligase SIZ1 (57), a key player in SA- and PAD4-mediated TIR-NBS-LRR signalling (58). Moreover, there is evidence for AMSHs regulating cell surface receptor proteins, involved in PTI, and intracellular NLR receptors involved in ETI in both plants and animals (21–24). The conservation of the Mp10 effector protein across the hemipteran lineage suggests that this is an important mechanism for plant immune-suppression, and likely has been a factor in the success of this group of insects.

Mp10 has evolved a function in plant-immune suppression from an ancestral group of proteins, the CSPs, with more established functions in perception of environmental chemicals (59, 60). CSPs, along with the odorant binding proteins (OBPs) and gustatory receptors (GRs) are arthropod-specific protein families that contribute to the chemosensory system of this group of organisms. CSPs and OBPs share structural similarities; both are small globular proteins formed from a series of α-helices stabilised by disulphide bonds and contain hydrophobic ligand-binding pockets that serve as binding sites for chemical signals (4). Whether the hydrophobic pocket of Mp10 contributes to its immune-suppressive function *in planta* remains unknown. Some evidence suggests that CSPs and OBPs may have a shared evolutionary origin, but this remains unclear (4). One sub-family of OBPs, the D7 and D7-like proteins in dipteran insects have evolved analogous functions to Mp10 - the D7 proteins are present in the saliva of mosquitos and inhibit blood clotting and inflammatory responses in the host animal (5, 61), and like Mp10 this function appears to be ancient and conserved amongst this group of insects.

## Materials and methods

### Plant materials and growth conditions

*N. benthamiana* WT seeds were provided by JIC Horticultural Service. *N. benthamiana* transgenic seeds expressing the N-terminus eGFP fusion of the calcium reporter aequorin under the 35S promoter (*35S:eGFP-aequorin*) were previously described (62). Seeds were sown directly in soil (Levington F2 Starter, SCOTTS) and incubated in a plant growth chamber (Hettich); one week after germination individual seedlings of approximately the same size were transplanted in 8-cm plastic rounded pots (one seedling per pot) containing about 0.4 l soil, and grown in a CER under a 16 h : 8 h, light/dark photoperiod at constant temperature of 22 °C and humidity of 80%. Plants were used for assays 5 or 6 weeks after sowing.

### Molecular cloning methods

All plasmids used in this study are listed in Supporting Information (S3 Table). For plasmids constructed using Gateway Cloning, coding sequences were amplified from cDNA using the primers listed in Supporting Information (Table S4), with attB1/attB2 adaptors. PCR products were cloned into the pDONR207 entry vector via BP Clonase (Thermo Fisher). Inserts were then transferred into the indicated destination plasmid by LR Clonase. For plasmids constructed using Golden Gate (GG) cloning, coding sequences were amplified using the primers indicated in Supporting Information (Table S4), and cloned into the pUAP-RFP by BpiI digestion-ligation. For Crispr cloning, primers for sgRNA sequences were annealed at 37 degrees, and cloned by BsaI digestion-ligation into pICSL002218A, which contains the sgRNA backbones, and the CAS9 expression cassette. Coding sequence modules were cloned into level one plasmids along with the corresponding components using BsaI digestion-ligation. For Gibson-assembly cloning, coding sequences were amplified using the primers indicated in Supporting Information (Table S4), and cloned into linearized plasmids (EcoRI-cut pLexAN or SfiI-cut pGADHA) using Gibson assembly master-mix (New England Biolabs).

### Agrobacterium-mediated transient gene expression

The binary vectors used for transient expression (listed in S3 table) were introduced into *A. tumefaciens* strain GV3101 containing the helper plasmid pMP90, which were subsequently infiltrated in leaves of 4- to 5-w-old *N. benthamiana* plants at a final concentration of OD_600_ = 0.3-0.5 in MMA buffer (10 mM MES pH = 5.5, 10 mM MgCl2, 100 µM acetosyringone). Leaf assays were conducted 2-3 days post infiltration.

### CRISPR-CAS9 mutagenesis of AMSH2

A CRISPR-associated transposon (CAST) including sgRNA-9 targeting the first AMSH2 exon, sgRNA-11 targeting the third AMSH2 exon, and 668 bp between the protospacer adjacent motif (PAM) sites, were cloned, together with the spCAS9 gene, in the pISCL02208 binary plasmid, which was transformed into *Agrobacterium tumefacians* GV3101 first and then into Arabidopsis Col-0 WT plants by the floral dip method (63). Kanamycin-resistant T1 transformants were screened for editing events by PCR with a primer pair flanking the two sgRNA sites that together amplify a 1477-bp product from WT, or a 760-bp product if a deletion is created between the two PAM sites. Multiple T1 independent lines showing the presence of a low molecular weight product consistent with a deletion between the two PAM sites were saved for seed collection. PCR products were Sanger sequenced, confirming the presence of the expected deletion.

### ROS burst assays

ROS production was determined using a luminol/peroxidase-based method as previously described (64, 65), with some modifications. For ROS burst, the youngest fully expanded leaves from 4- to 5-w-old *N. benthamiana* WT plants were infiltrated with *A. tumefaciens* carrying the respective constructs (i.e., eGFP-alone, eGFP-Mp10/MpCSP4, eGFP-AgCSP4, eGFP-ApCSP4, eGFP-BtCSP4, eGFP-CtCSP4, or eGFP-DmCSP4). Two days after agroinfiltration, at least 16 leaf discs of 4 mm diameter were obtained from the agroinfiltrated patches of at least three plants and incubated in white 96-well polystyrene microplates (Grenier Bio-One LUMITRAC^TM^) containing 200 μl of ultrapure water per well, and covered with aluminium foil. After 1 day, the water was replaced with 100 μl water solution containing 10 μg/mL horseradish peroxidase (Sigma-Aldrich) and 17 μg/mL luminol (Sigma-Aldrich) supplemented with 100 nM flg22 peptide (QRLSTGSRINSAKDDAAGLQIA, EZBiolab). Change of luminescence was measured every 30 s for at least 1 hour using a Photek camera (East Sussex, UK) and the IFS32 imaging software (Photek).

### Ca^2+^ burst assays

*N. benthamiana* expressing the aequorin calcium sensor (62) were grown and agroinfiltrated as for ROS bursts. Leaf-discs were collected and incubated overnight in the dark in 12.5 μM Coelenterazine (Biosynth) in 96-well plates. Coelenterazine solution was exchanged for 100 ul 100 nM flg22 and plates were imaged using a Photek photon counting camera as described for ROS bursts. Data were collected for 30 mins post elicitation.

### Enrichment of membrane-associated proteins

Enrichment of membrane proteins was performed from *N. benthamiana* agroinfiltrated leaves for the purpose of conducting co-immunoprecipitation assays, as previously described (29). Briefly, each sample containing different combinations of constructs used for the GFP- and Mp10-pull downs were infiltrated in one entire leaf per plant. After 3 DPI, the infiltrated leaves (approximately 1.6 g each) were detached, snap-frozen in liquid nitrogen, and ground to a fine powder using mortars and pestles. The powder was then supplemented with 16 mg polyvinylpolypyrrolidone (PVPP) and then homogenized in 1.6 ml of protein extraction buffer (100 mM Tris-HCl at pH 8.8, 150 mM NaCl, 1 mM EDTA, 10% glycerol, 20 mM NaF, 1 mM PMSF, 1X Sigma-Aldrich protease inhibitor cocktail, 1 mM Na_2_MoO_4_, 50 mM β-glycerophosphate, 10 mM Na_3_VO_4_). Then, cell lysates containing total proteins (soluble plus membrane-associated proteins) were cleared twice by centrifugation at 5,000 g for 5 min at 4 °C. Membrane-associated protein fractions were separated from cytoplasmic soluble proteins by ultracentrifugation at 56,000 rpm for 36 min at 4 °C, using an Optima^TM^ MAX-E ultracentrifuge (Beckman Coulter) equipped with a TLA120.2 fixed angle rotor (Beckman Coulter) and 1 mL polypropylene tubes (Open-Top Thickwall, Beckman Coulter). The pellets (containing the membrane fractions and membrane-associated proteins) were subsequently solubilized in 500 μl of membrane protein solubilization buffer (100 mM Tris-HCl pH 7.3, 150 mM NaCl, 1 mM EDTA, 10% glycerol, 20 mM NaF, 1% Triton X-100, 1 mM PMSF, 1X Sigma-Aldrich protease inhibitor cocktail) with sonication, using a Soniprep 150 ultrasonic disintegrator (4% amplitude, 5s:5s ON:OFF cycle for about 10 cycles or until completely dissolved). To remove any insoluble particles, resuspended membrane fractions were ultra-centrifugated as described in the previous ultracentrifugation step. Cleared supernatants were subjected to pull down assays.

### GFP- and Mp10-pull downs in tobacco leaves

Pull downs of GFP-tagged proteins were performed using GFP-Trap Magnetic Agarose (chromotek), according to manufacturer’s instruction. Briefly, membrane-associated protein enriched fractions were diluted in dilution buffer (10 mM Tris-HCl pH 7.5, 150 mM NaCl, 0.5 mM EDTA) at 2 : 3 (proteins : buffer) ratio and incubated with GFP magnetic beads for over-night at 4 °C with rotation. Proteins were eluted at 95 °C in 60 μl of loading buffer (2X Novex™ NuPAGE™ LDS Sample Buffer (Invitrogen™) supplemented with 10% β-mercaptoethanol) and subjected to western blot and mass spectrometry analyses.

Pull downs of Mp10 were performed using Pierce^TM^ Co-Immunoprecipitation (Co-IP) Kit (Thermo Scientific^TM^), according to manufacturer’s instruction. Anti-Mp10 antibody (6) was coupled to resin and incubated with membrane-associated protein enriched fractions in IP wash buffer (25 mM Tris-HCl pH 7.4, 150 mM NaCl, 1 mM EDTA, 1% NP-40, 5% glycerol) at 2 : 1 (proteins : buffer) ratio over-night at 4 °C with rotation. Proteins were eluted in 50 μl of Elution Buffer (pH 2.8, containing primary amine) and subjected to western blot and mass spectrometry analyses.

### Western blot analyses

Proteins in loading buffer were resolved on 4-12% Mini-PROTEAN® TGX™ Precast Gels (BIO-RAD) in 1x Tris/Glycine/SDS Buffer (BIO-RAD) and transferred to 0.2 μm PVDF or nitrocellulose membrane using Trans-Blot Turbo Transfer System (BIO-RAD). The membranes were blocked in blocking buffer made of washing buffer (1x PBS, 0.2% Tween 20) supplemented with 10% non-fat dry milk and 5% BSA, and probed with primary antibodies (diluted in blocking buffer), as follows: anti-GFP-HRP (1 : 4000, Santa Cruz); anti-RFP-HRP (1 : 2500, Santa Cruz); anti-Flag-mouse (1 : 2500, Agrisera); anti-HA-mouse (1 : 2000, Sigma-Aldrich); anti-Mp10-chicken (1 : 2500) (6); or anti-GST-rabbit (1: 4000, Sigma-Aldrich). This was often followed by incubations with secondary antibodies (diluted in blocking buffer), as follows: anti-mouse-HRP (1 : 20000, Sigma-Aldrich); anti-chicken-HRP (1 : 20000, Sigma-Aldrich); or anti-rabbit-HRP (1 : 20000, Sigma-Aldrich). HRP presence was detected via incubation with ECL™ detection reagents (Millipore). The subsequent chemiluminescence signal was detected with Amersham ImageQuant 800 (Cytiva) and analysed with Amersham IQ800 Control software (v.1.2.0).

### Mass spectrometry analyses

For MS analyses, 50 ul of Co-IP sample were mixed with 300 ul of chloroform : methanol, 1 : 4 solution, kept on ice for 30 min, and centrifugated at high speed (14000 rpm) for 15 minutes at 4 °C. Protein pellets were washed two times with 1 ml methanol and one time with 1 ml acetone.

For protein identification by mass spectrometry, we used a nanoLC-MS/MS on an Orbitrap EclipseTM TribridTM mass spectrometer coupled to an UltiMate® 3000 RSLCnano LC system (Thermo Fisher Scientific, Hemel Hempstead, UK). The predicted *N. benthamiana* proteome from the Nbe_v1.1 genome assembly (https://nbenthamiana.jp/) (66) was used to identify peptides and assign them to proteins along with their functional descriptions. Data of combined datasets or independent replicates were analysed using the Thermo Scientific Proteome Discoverer 3.0 software (PD3.0). Abundance values derived from the mass spectrometry readings were normalized to account for unequal total protein concentration in each sample (with imputation). Next, treatments 1) eGFP-AMSH2 + Flag-Mp10, 2) eGFP-alone + Flag-Mp10, 3) eGFP-AMSH2 + Flag-alone, 4) eGFP-alone + Flag-alone were compared by abundance ratios between treatments 1 *vs* 2 and 3 *vs* 4 for the eGFP-AMSH2 co-immunoprecipitation experiments, and 1 *vs* 3 and 2 *vs* 4 for the Mp10 co-immunoprecipitation experiments. This allowed for the identification of candidate proteins that co-immunoprecipitated with eGFP-AMSH2 in the presence and absence of Mp10 and with Flag-Mp10 in the presence and absence of eGFP-AMSH2. A background-based t-test in PD3.0 differentiated candidate proteins from the majority of proteins, with p-values adjustments as described (67), and a threshold of at least 4-fold change combined with the adjusted p-value significance threshold of ≤0.05 was applied. In addition, proteins with a non-adjusted p-value threshold of <0.05 were also inspected for the presence of groups of proteins with shared biological functions. These remaining candidate proteins were then analysed to inform possible mechanisms of PTI suppression by the Mp10-AMSH2 interaction and to propose a normal function of AMSH2. A more detailed description of the mass spectrometry methods is available in Supporting Information (S1 Protocol). The mass spectrometry proteomics data were deposited to the ProteomeXchange Consortium (68) via the PRIDE (69) partner repository with the dataset identifier PXD055941 and 10.6019/PXD055941 (35).

### FLIM-FRET assays

For FLIM-FRET assays, eGFP-tagged Mp10, or eGFP-alone, were co-expressed with mCherry-tagged AMSH proteins, or mCherry fused to aquaeorin in *N. benthamiana* leaves via *A. tumefaciens*-mediated agroinfiltration as described above (in agrobacterium-mediated transient gene expression methods). Abaxial epidermal cells of leaf sections were imaged 2-3 days after infiltration using a Leica Stellaris 8 microscope. Images were captured detecting fluorescence from eGFP (laser, ex. 488 nm, em. 509-534 nm) mCherry (laser, ex. 587 nm, em. 603-625 nm) and chlorophyll (laser, ex. 587 nm, em 687-712 nm). To avoid bleed through of chlorophyll fluorescence into the eGFP channel, regions lacking chlorophyll but showing green and red fluorescence, indicating the presence of eGFP- and mCherry-tagged proteins, were selected for FLIM analysis. Fluorescence lifetime data derived from eGFP were collected from these regions in FLIM mode (ex. 488 nm, em. 525-530 nm) at 128×128 resolution until 1000 photons per pixel at the most intense regions of the image. Instrument response function was captured using erythrosine on each day of data collection. FLIM data were analysed using Leica Application Suite X (LAS X) FLIM FCS software. Fluorescence lifetime decay curves of eGFP-alone control samples were modelled as a 2-component exponential function, and all samples from each experimental set were modelled against the fluorescence lifetime from the corresponding control samples to derive values for fluorescent lifetime and %FRET efficiency for each image collected. Differences in lifetime and FRET were determined by a one-way ANOVA with a Tukey HSD test. Percentages (%) of FRET efficiency were mapped to the images (as shown in Figs 2F, 2H, S3B, S3D and S3F), and phasor plots were generated for whole images as well as for regions of interest (ROI) with the highest and lowest FRET efficiency, showing that the FRET signal was associated with a clockwise shift on the phasor plot consistent with bona-fide FRET (31). Full collection of image and data collection used for these experiments, including metadata, are deposited in Zenodo (70).

### Confocal microscopy

Confocal microscopy analysis was performed on a Leica TCS SP8X confocal DM6 microscope with a 63x water-immersion objective, using Leica Application Suite X (LAS X) software (3.5.7.23225). Excitations of eGFP and chlorophyll signals were achieved with a 488 nm Argon laser with emission, respectively, at 495–545 nm and 690-710 nm. Excitations of RFP signal was achieved with a 590 nm white light laser (WLL) with emission at 605–650 nm. Full collection of image and data collection used for these experiments, including metadata, are deposited in Zenodo (70).

### Heterologous protein production in *E. coli*

Plasmids used for protein expression and purification (listed in S3 Table) were transformed into *Escherichia coli* Rosetta (DE3) cells (Merck Chemicals). A single positive colony of fresh *E. coli* was grown up to OD600 ∼ 0.5 in 1 L Terrific Broth medium (24 g/L yeast extract, 20 g/L tryptone, 0.4% glycerol, phosphate buffer (0.017 M KH_2_PO_4_, 0.072 M K_2_HPO_4_)), and then induced with 200 μM IPTG at 16 °C for 16 h.

### GST-pull downs from *E. coli*

GST-pull downs were performed from Rosetta (DE3) *E. coli* cells co-transformed with i) GST-GBP (71), GST-AMSH1, GST-AMSH2, or GST-AMSH3 (32) and ii) His-Mp10 (or without His-Mp10, as control), as previously described (72). Briefly, bacterial pellets were re-suspended in 50 ml buffer A (50 mM Tris-Hcl pH 7.5, 300 mM NaCl, 10% glycerol, 0.2% Triton X-100, 1x Roche cOmplete EDTA-free protease inhibitor), sonicated (time, 5 m; cycle, 10 s : 5 s, ON : OFF; amplitude 40%) and centrifugated (15000 x*g*; 4 °C). Cleared supernatants were incubated with Glutathione Sepharose 4B (75 μl bed volume) for 2 h at 4 °C with rotation. Proteins were eluted at room temperature in 100 μl of 40 mM reduced glutathione and subjected to western blot analysis, as described above.

### Protein purification in *E. coli* and *in vitro* gel filtration assays

Purification of 6xHis-MBP-3C-Mp10 or 6xHis-GB1-3C-AMSH2 from bacteria was performed using ÄKTA go™ protein purification system (Cytiva). Briefly, bacterial pellets were re-suspended in 50 ml immobilized metal ion affinity chromatography (IMAC) lysis buffer (50 mM Tris-HCl pH 8.0, 500 mM NaCl, 50 mM glycine, 5% glycerol, 20 mM imidazole), sonicated (time, 5 m; cycle, 10 s : 5 s, ON : OFF; amplitude 40%) and centrifugated (18000 rpm; 4 °C). Clear supernatants were flowed into Ni-charged HisTrap™ High Performance 5 ml column (Cytiva), eluted in IMAC elution buffer (50 mM Tris-HCl pH 8.0, 500 mM NaCl, 50 mM glycine, 5% glycerol, 500 mM imidazole). Ni-purified protein was loaded into HiLoad® 26/600 Superdex® 200 pg preparative gel filtration chromatography column (Cytiva) and eluted in gel filtration buffer (20 mM HEPES pH 7.5, 150 mM NaCl) supplemented with 1 mM TCEP (only for AMSH2 purification). To remove the tag, the protein was incubated with 100 μg His-tagged HRV 3C protease for overnight at 4 °C. To separate cleaved tag and protease from the target protein, a reverse-IMAC purification (i.e., the flow-through is retained) was performed. Finally, the target protein was further purified by preparative gel filtration chromatography, as described above. For testing complex formation, equal amount (molecular weight) of purified Mp10 and AMSH2 were mixed in gel filtration buffer, incubated on ice for 30 mins, and subjected to analytical gel filtration chromatography.

### *In vitro* DUB activity assays

DUB activity assays were performed as previously described (32, 72). Briefly, 1 μg K63-linked di-ubiquitin substrate (ENZO LIFE SCIENCES UK LTD) was incubated at 30 °C for the indicated times in 10 μl DUB buffer (50 mM Tris-HCl pH 7.2, 25 mM KCl, 5 mM MgCl_2_, 0.1 mM DTT) ± 1 μg AMSH2. To examine the Mp10 effect on AMSH2 activity, AMSH2 was preincubated with 1.4 μg Mp10 for 30 min at 4 °C before addition of di-ubiquitin substrate. Reactions were terminated at 4 °C. Proteins were diluted with loading buffer (1X Novex™ NuPAGE™ LDS Sample Buffer (Invitrogen™) supplemented with 1% β-mercaptoethanol), boiled for 5 min, resolved on 4-12% or 16% Mini-PROTEAN® TGX™ Precast Gels (BIO-RAD), and stained with Coomassie (ReadyBlue® protein gel stain) or Silver staining (Pierce™ Silver Stain Kit).

For fluorescence-based DUB assay, protein samples containing 1) AMSH2, 2) AMSH2 + Mp10, or 3) gel filtration buffer (negative control), were incubated in 100 μl TAMRA DUB buffer (50 nM K63-linked FRET TAMRA Position 3, 50 mM Tris-HCl pH 7.5, 100 mM NaCl, 0.1% Pluronic F-127, 0.5 mM TCEP) in 96-well black plate. Change of fluorescence was measured every minute over a period of 1 h using a VarioSkan plate reader (ex. 530 nm; em. 590 nm; 23 °C).

### Y2H screen and Y2H assays

A yeast-2-hybrid library was constructed using combined mRNAs isolated from non-infected *Arabidopsis thaliana* Col-0 plants and *A. thaliana* Col-0 plants exposed to the aphid species *Myzus persicae* (peach-potato aphid) or *Acyrthosiphon pisum* (pea aphid), leafhopper species *Macrosteles quadrilineatus* (aster leafhopper) or *Dalbulus maidis* (corn leafhopper), or *A. thaliana* Col-0 plants infected with the Aster Yellows Witches Broom (AY-WB) phytoplasma (*Candidatus* Phytoplasma asteris). The cDNAs were cloned into the SfiI site of pGAD-HA prey vector (Dualsystems). For screening the library, the sequence corresponding to the mature part of Mp10 (without signal peptide) was cloned into the EcoR1 site of the bait vector pLexA-N and transformed into yeast (*Saccharomyces cerevisiae* NMY51). Minimal autoactivation of the Mp10 plasmid was found on media lacking histidine and supplemented with 2 mM 3-amino triazole (3-AT). The Mp10 bait vector was co-transformed with the *A. thaliana* Col-0 library into NMY51 according to the manufacturer’s instructions (Dualsystems), screening approximately 5 x 10^6^ colonies on the above media, resulting in 6 colonies that showed growth. The prey plasmids were isolated from these 6 colonies and tested for self-activation by co-transforming into NMY51 with the empty pLexA-N bait vector. Three colonies that did not show self-activation were sanger sequenced and all contained the full length AMSH2 coding sequence.

Homologues of Mp10 (without signal peptide) from other hemipteran insects were similarly cloned into the EcoRI site of pLexaN, and full-length coding sequences of AMSH proteins cloned into the SfiI site of pGAD-HA. Pairwise yeast-2-hybrid experiments were performed by co-transformation of pairs of bait- and prey-plasmids into NMY51, as per the manufacturer’s instructions, and screened on media lacking histidine and/or adenine and a range of concentrations of 3-AT.

### Bioinformatics and phylogenetic analyses

The phylogenetic analysis of CSPs from different hemipterans showed in Fig 1A is adapted from our previously published preprint manuscript (25). To identify *M. persicae* CSPs, published pea and cotton/melon aphid CSPs (73, 74) were BLASTP searched against the GPA clone O genome database. Hits with e<10*^-5^* were reciprocally BlastP searched against the *A. pisum* genome, and those with an annotated CSP as the top hit were kept for further analysis. CSPs were also identified in the sequenced genomes of *A. pisum*, *D. noxia* (Russian wheat aphid), *N. lugens* (rice brown planthopper), *C. lectularius* (the bedbug) and *R. prolixus* (the kissing bug). Moreover, the transcriptomes of *B. brassicae* (cabbage aphid), *C. tenellus* (beet leafhopper), *D. maidis*, *M. quadrilliniatus* and *B. tabaci* (tobacco whitefly) were sequenced using RNAseq (NCBI SRA PRJNA318847-PRJNA318851). Coding sequences (CDS) were predicted from the *de novo* assembled transcriptomes and CSPs were identified in all sets of predicted protein sequences based on reciprocal best blast hits to the annotated set of *M. persicae* CSPs. Annotated CSP sequences from the genome and transcriptome data were aligned and phylogenetic analysis was carried out using this alignment. Full details of the bioinformatics and phylogenetic analysis carried out can be found in our previously published preprint manuscript (25), SI Appendix, Materials and Methods.

## Acknowledgements

We thank Erika Isono (University of Konstanz), Sophien Kamoun, Mark Youles and Freddy Boutrot (The Sainsbury Laboratory, Norwich), Tolga Bozkurt (Imperial College London), Silke Robatzek (LMU Munich Biocenter), Cyril Zipfel (University of Zurich), and Kazuhisa Nakayama (Kyoto University) for providing plasmids, other materials, and valuable discussions. We also thank members of the Hogenhout lab, especially Friederike Bernsdorff, Federico-Gabriel Mirkin, Qun Liu, and Yeshveer Singh, for their insights. We acknowledge the John Innes Centre (JIC) Horticultural Services for plant cultivation, the JIC Bioimaging Platform for training and support in confocal microscopy and FRET-FLIM, the JIC Insectary and Entomology Facility for insect rearing, and the TSL Synbio service for making, preparing and/or supplying plasmids. This work was funded by UKRI Biotechnology and Biological Sciences Research Council (BBSRC) grants to SAH (BB/V008544/1 and BB/N009169/1), BBSRC Doctoral Training Partnership fellowships to CD, JC, and CW, and a Sainsbury PhD Studentship in Plant Science to JJ. Additional support was provided by BBSRC Institute Strategy Programmes (BBS/E/J/000PR9797 and BBS/E/JI/230001B) awarded to the John Innes Centre, which is grant-aided by the John Innes Foundation.

## Author contributions

Conceptualization, MG, SAH, STM; Data Curation, All; Formal Analysis, All; Funding Acquisition, SAH; Investigation, All; Methodology, All; Project administration, SAH; Resources, CD, MG, SAH, JJ, AK, TCM, STM, CW; Supervision, MG, SAH, STM; Validation, All; Visualization, JC, CD, MG, JJ, AK, NK, TCM, STM, CW; Writing – original draft preparation, MG, SAH, STM; Writing – Review and Editing, All.

## Data availability

All data is included in the supplementary information, or associated database repositories (PRIDE, Zenodo accessions PXD055941, zenodo.13752241, respectively).

## Supporting information

### Other Supporting information materials for this manuscript include the following

Tables S1 and S2 (provided as separate files)

**S1 Fig.**
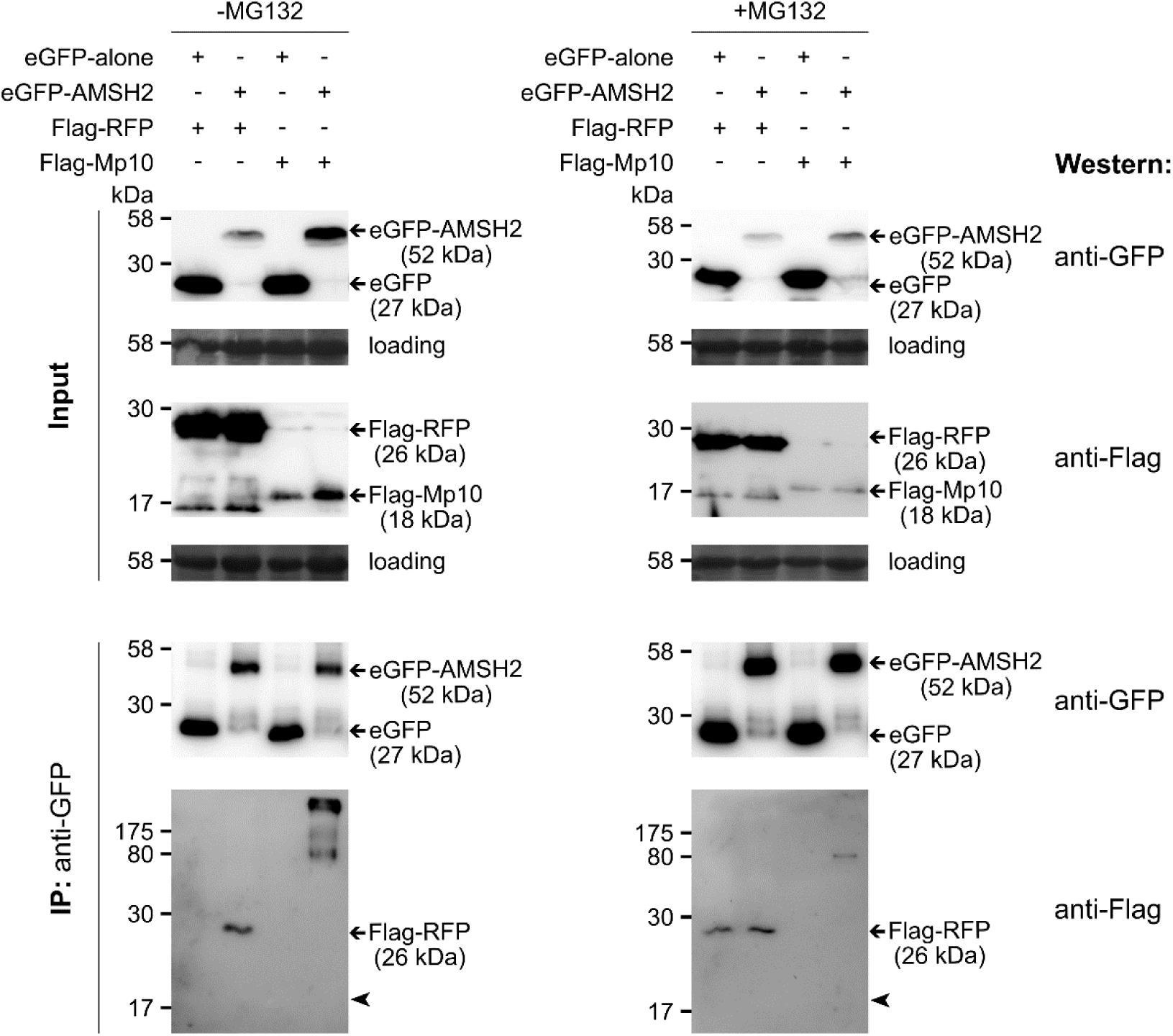
GFP co-immunoprecipitation of eGFP-AMSH2 with Flag-Mp10 from total soluble protein extracts. Co-immunoprecipitation (co-IP) were conducted from *N. benthamiana* leaves transiently expressing Flag-Mp10 or Flag-RFP, as control, with eGFP-alone (control) or eGFP tagged Arabidopsis AMSH2, in presence/absence of the proteasome inhibitor MG132, as indicated, followed by western blot analyses using antibodies to eGFP and Flag. Protein sizes indicated by molecular weight markers (kDa). Expected bands marked with arrows; arrowheads denote expected location of Flag-Mp10. Loading visualized with amido black staining.

**S2 Fig.**
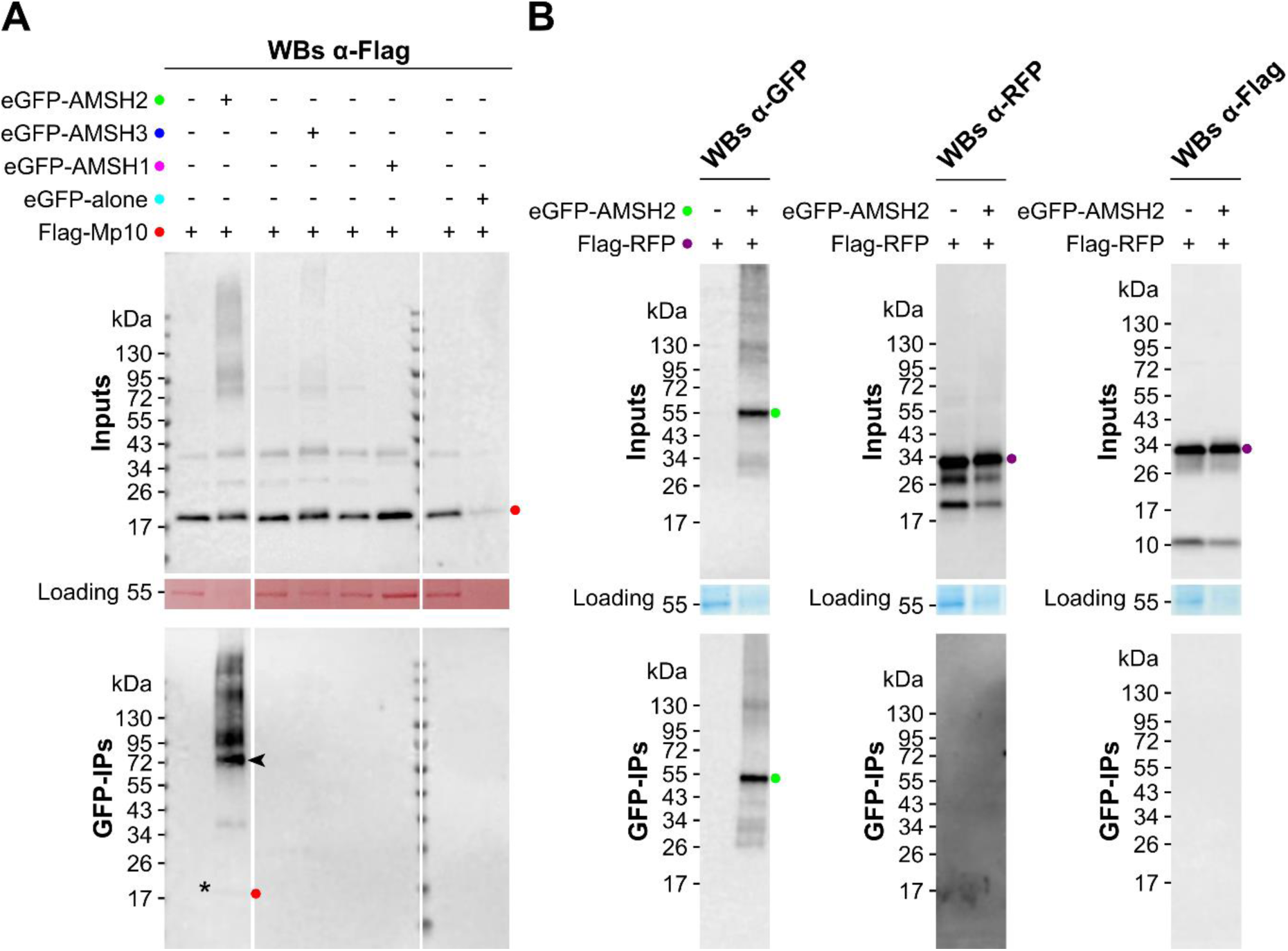
GFP co-immunoprecipitation of eGFP-AMSH2 with Flag-Mp10 from membrane-enriched protein extracts. AMSH2 pulls down Mp10 from *N. benthamiana* membrane fractions. Co-immunoprecipitation (co-IP) were conducted from *N. benthamiana* leaves transiently producing (**A**) Flag-Mp10 or (**B**) Flag-RFP (control) with eGFP-alone (control) or eGFP tagged Arabidopsis AMSH1, AMSH2, or AMSH3, as indicated, followed by western blot analyses using antibodies to Flag (A-B), and eGFP and RFP (B). Protein sizes indicated by molecular weight markers (kDa). Expected bands marked with colored dots; asterisk denotes Flag-Mp10 pulled down with AMSH2; arrowheads denote a putative modified form of Mp10, and/or an oligomeric state of Mp10, or a higher-order complex containing the Mp10 protein that is apparently resistant to denaturation by SDS-PAGE. Loading visualized with Ponceau S or Coomassie Blue staining.

**S3 Fig.**
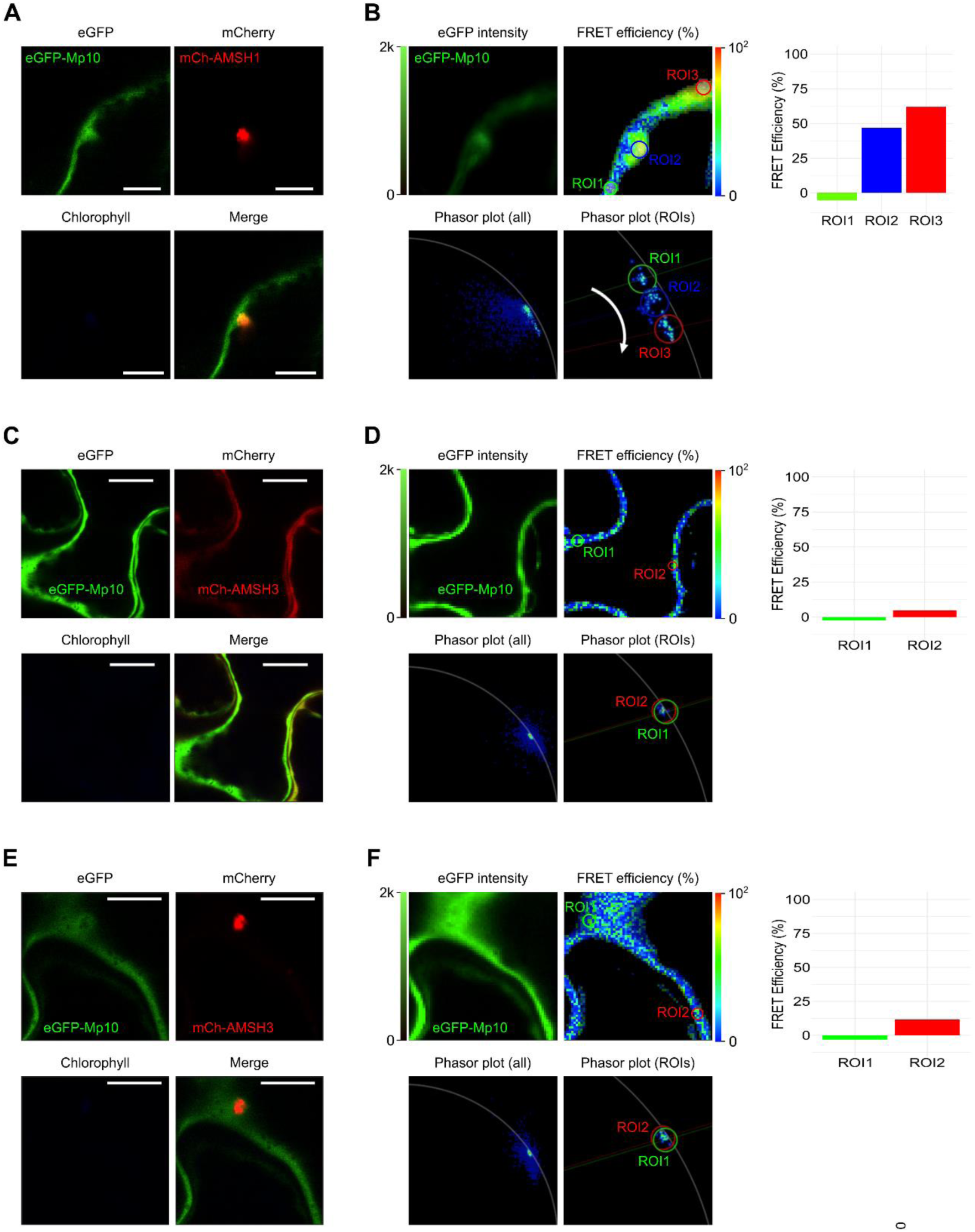
eGFP-Mp10 interacts with mCherry-AMSH1, but not with mCherry-AMSH3, in FLIM-FRET assays. FRET analysis of eGFP-Mp10 in presence of mCherry-AMSH1 (**A, B**) or mCherry-AMSH3 (**C, D**). In A, C, E, confocal images show subcellular localization of eGFP-Mp10 with mCherry-AMSH1 or mCherry-AMSH3. In B, D, F, panels show eGFP-Mp10 intensity (upper left), FRET efficiency heat maps (upper right), and phasor plots for full images (lower left) and selected regions of interest (ROIs) (lower right). Graphs to the right show the extracted FRET efficiency values (y-axis) for the selected ROIs (x-axis).

**S4 Fig.**
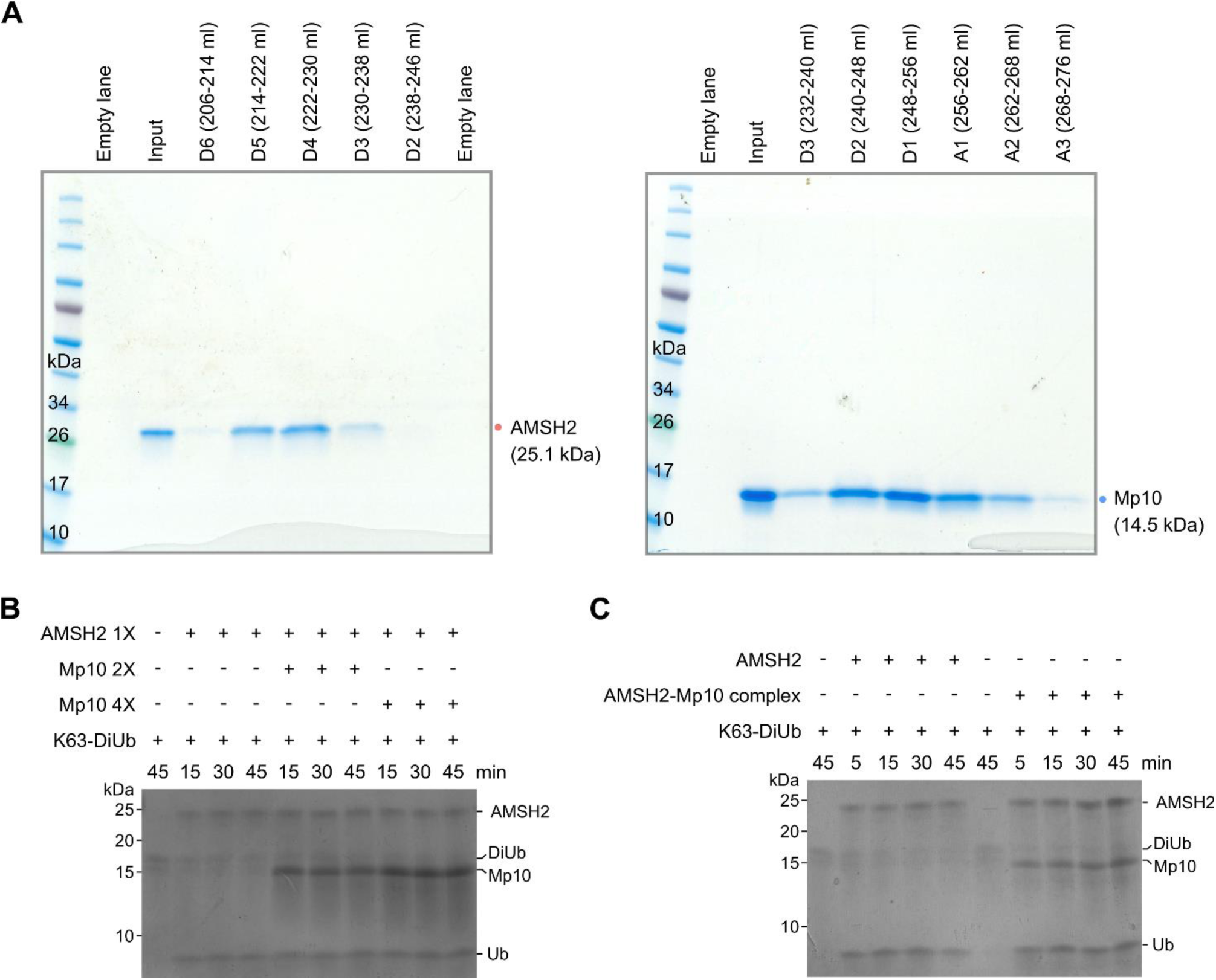
Purification of AMSH-Mp10 protein complexes and DUB activity assays. (**A**) Coomassie-stained SDS-PAGE gels of fractions collected from preparative gel filtration column at various elution volumes indicate recombinant protein sizes with colored dots. The collected fractions were pooled and concentrated for subsequent intact mass spec and analytical gel filtration analyses (see Fig 3B and 3C). (**B-C**) *In vitro* de-ubiquitination (DUB) assays using unlabeled K63-linked di-ubiquitin substrate in presence/absence of AMSH2 alone or AMSH2 with Mp10 at molar ratios 1:2 or 1:4 (B) and in presence of AMSH2 alone or AMSH2-Mp10 complex at equimolar concentration (C). Silver-stained 16% SDS-PAGE gels show reactions at specified time points, with MW markers on the left and expected bands on the right.

**S5 Fig.**
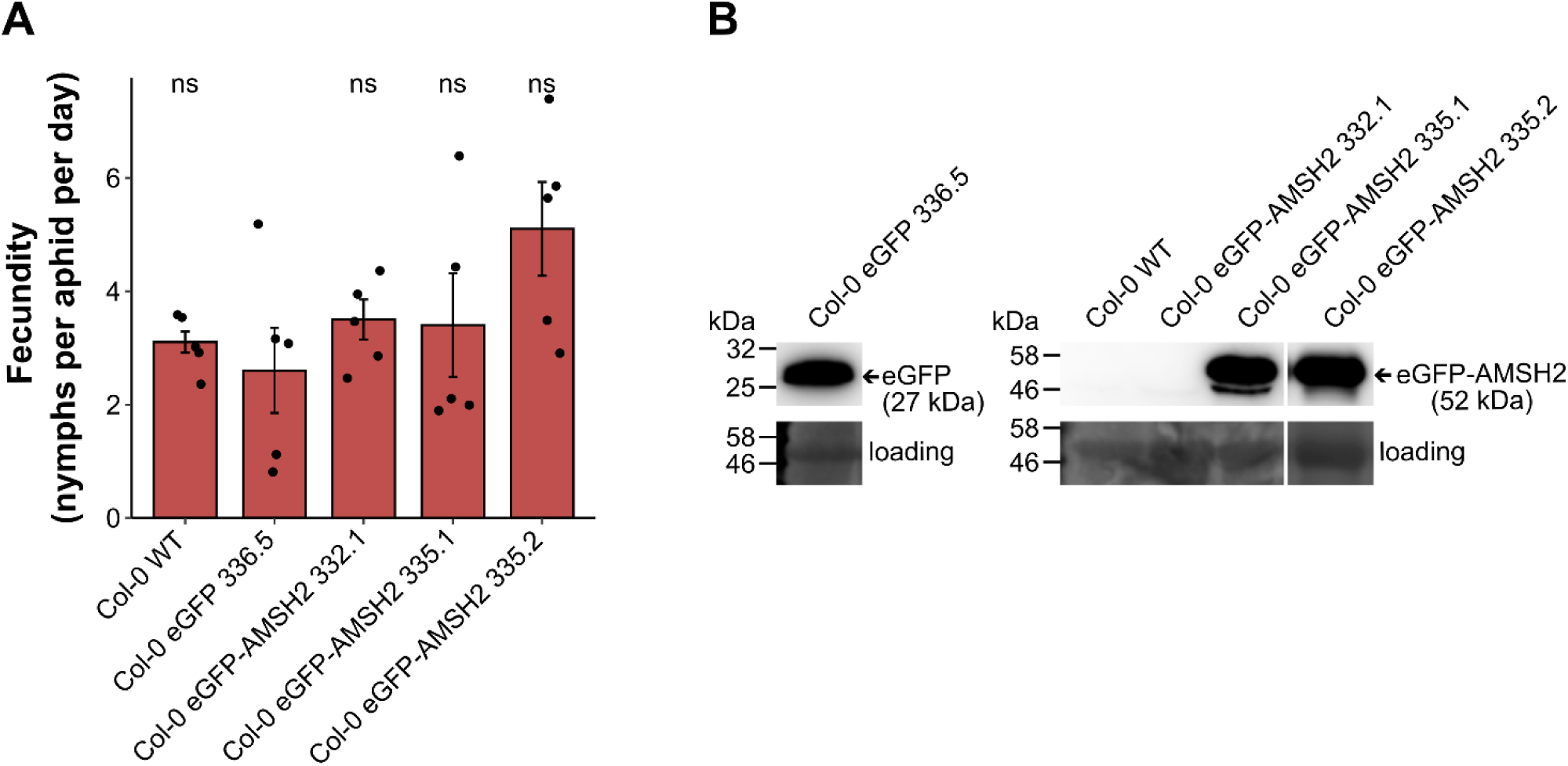
Aphid fecundity is not affected in Arabidopsis transgenic lines overexpressing AMSH2. (**A**) Aphid fecundity was not altered in three independent Arabidopsis Col-0 transgenic lines overexpressing eGFP-AMSH2, and in Col-0 WT, compared to transgenic lines overexpressing eGFP-alone. Y-axis: number of nymphs produced per mother aphid per day; x-axis: genotype, Col-0 WT, Col-0 eGFP-alone, Col-0 eGFP-AMSH2 (three independent lines, as indicated). Bar-plots show average ± standard error (SE) and observations as black filled circles; ns, not significant compared to Col-0 eGFP (control) according to ANOVA with Tukey HSD test. (**B**) Protein expression verification for fecundity assay. Western blot detects eGFP in Arabidopsis Col-0 plants stably overexpressing eGFP-alone or eGFP-AMSH2 (three independent lines, as indicated). MW markers are indicated on the left; expected bands are marked by arrows. Loading visualized with Ponceau S staining.

**S6 Fig.**
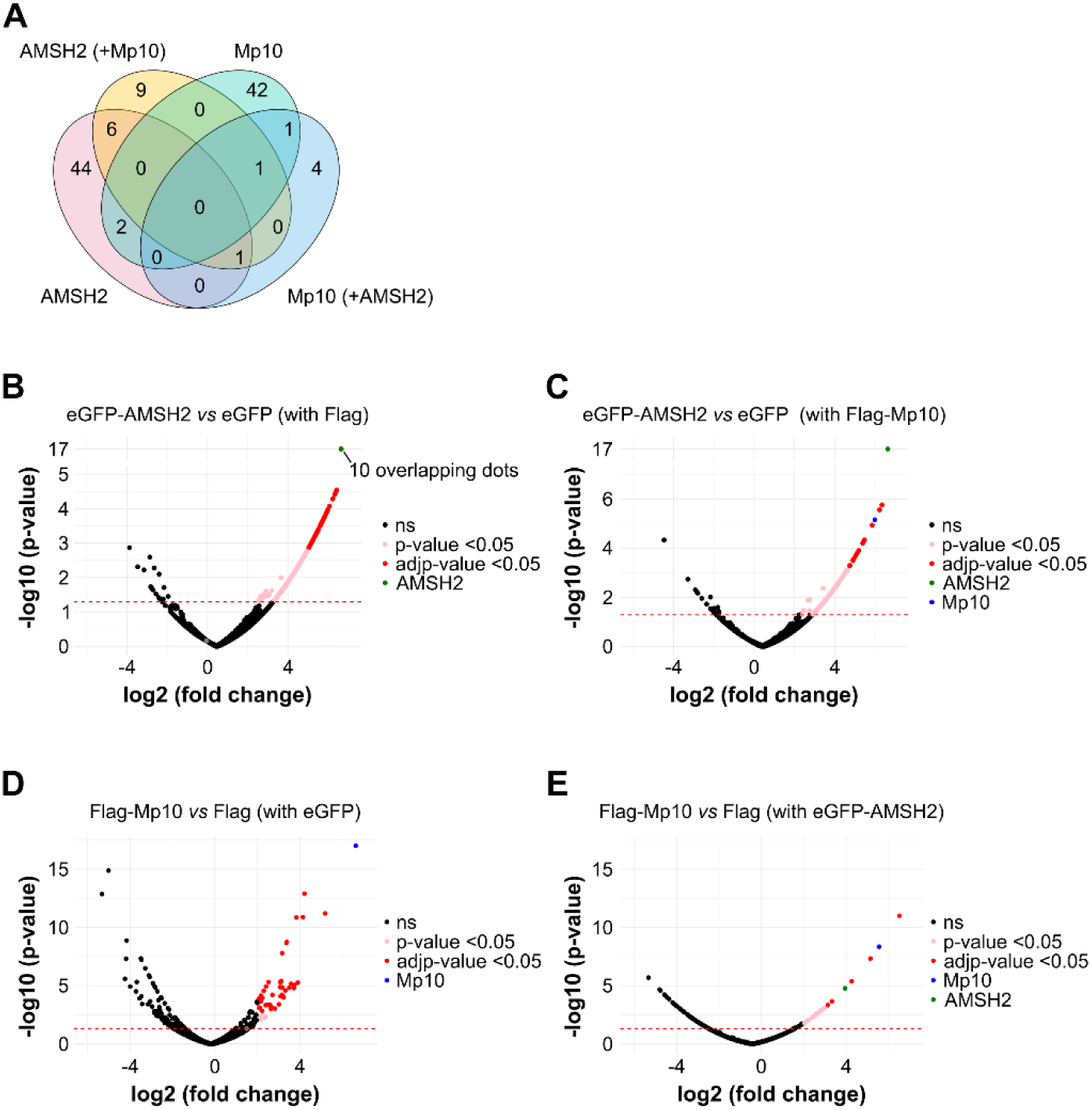
Identification of Mp10-AMSH2 interactors using co-IP-MS/MS in *N. benthamiana*. (**A**) Venn diagram showing overlap of interacting proteins for (i) eGFP-AMSH2, (ii) eGFP-AMSH2 + Flag-Mp10, (iii) Flag-Mp10, and (iv) Flag-Mp10 + eGFP-AMSH2 from co-IP-MS/MS datasets (fold change > 4; adjusted p-value < 0.05). (**B-E**) Volcano plots showing co-IP-MS/MS results from *N. benthamiana*, as follows: (A) eGFP-AMSH2 + Flag-alone vs. eGFP-alone + Flag-alone (control); (C) eGFP-AMSH2 + Flag-Mp10 vs. eGFP-alone + Flag-Mp10 (control); (C) Flag-Mp10 + eGFP-alone vs. Flag-alone + eGFP-alone (control); (D) Flag-Mp10 + eGFP-AMSH2 vs. Flag-alone + eGFP-AMSH2 (control). In each Volcano plot, interacting proteins in control groups are shown on the left, and those in experimental groups on the right. Statistically significant interactors (with fold change > 4) are highlighted as red dots (adjusted p-value < 0.05) and pink dots (p-value < 0.05) above the threshold line (red dotted line), while non-significant (ns) interactors appear as black dots. The eGFP-AMSH2 protein is marked as a green dot, and the Flag-Mp10 protein as a blue dot. Source data are available via ProteomeXchange with identifier PXD055941.

**S7 Fig.**
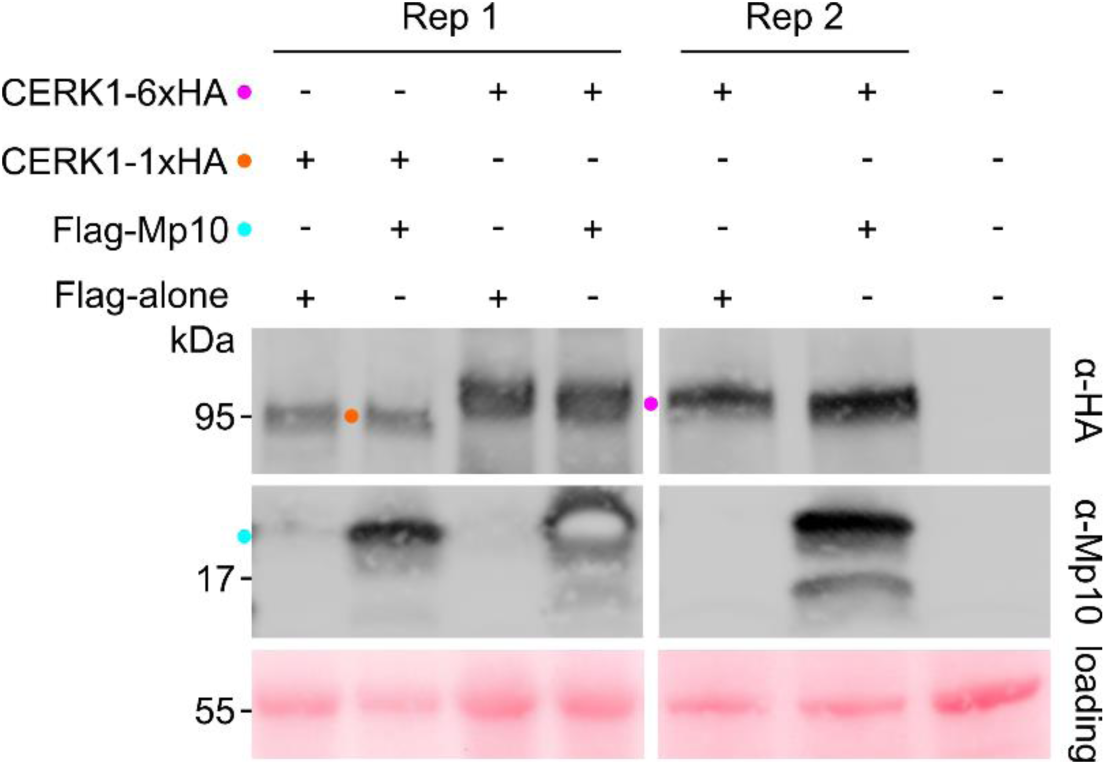
Mp10 does not affect the abundance of CERK1 cell-surface receptor-like kinase. Western blot analyses of Arabidopsis CERK1-1xHA and CERK1-6xHA protein stabilization in presence/absence of Flag-Mp10 using antibodies to HA (top panel) or Mp10 (middle panel). Protein sizes indicated by molecular weight markers (kDa). Expected bands marked with colored dots. Loading visualized with Ponceau S staining.

**S1 Table.** Selected proteins identified with LC-MS/MS that co-immunoprecipitated with Flag-Mp10 in presence or absence of eGFP-AMSH2. Table shows the average abundance ratio across three biological replicates of proteins Co-IP’d with Mp10 in the presence (Ratio 1/3) or absence (Ratio 2/4) of eGFP-AMSH2, together with the p-value and adjusted p-value, together with details of peptide abundance and confidence. Source data are available via ProteomeXchange with identifier PXD055941.

**S2 Table.** Selected proteins identified with LC-MS/MS that co-immunoprecipitated with eGFP-AMSH2 in presence or absence of Flag-Mp10. Table shows the average abundance ratio across three biological replicates of proteins Co-IP’d with eGFP-AMSH2 in the presence (Ratio 1/2) or absence (Ratio 3/4) of Mp10, together with the p-value and adjusted p-value, together with details of peptide abundance and confidence. Source data are available via ProteomeXchange with identifier PXD055941.

**S3 Table.**
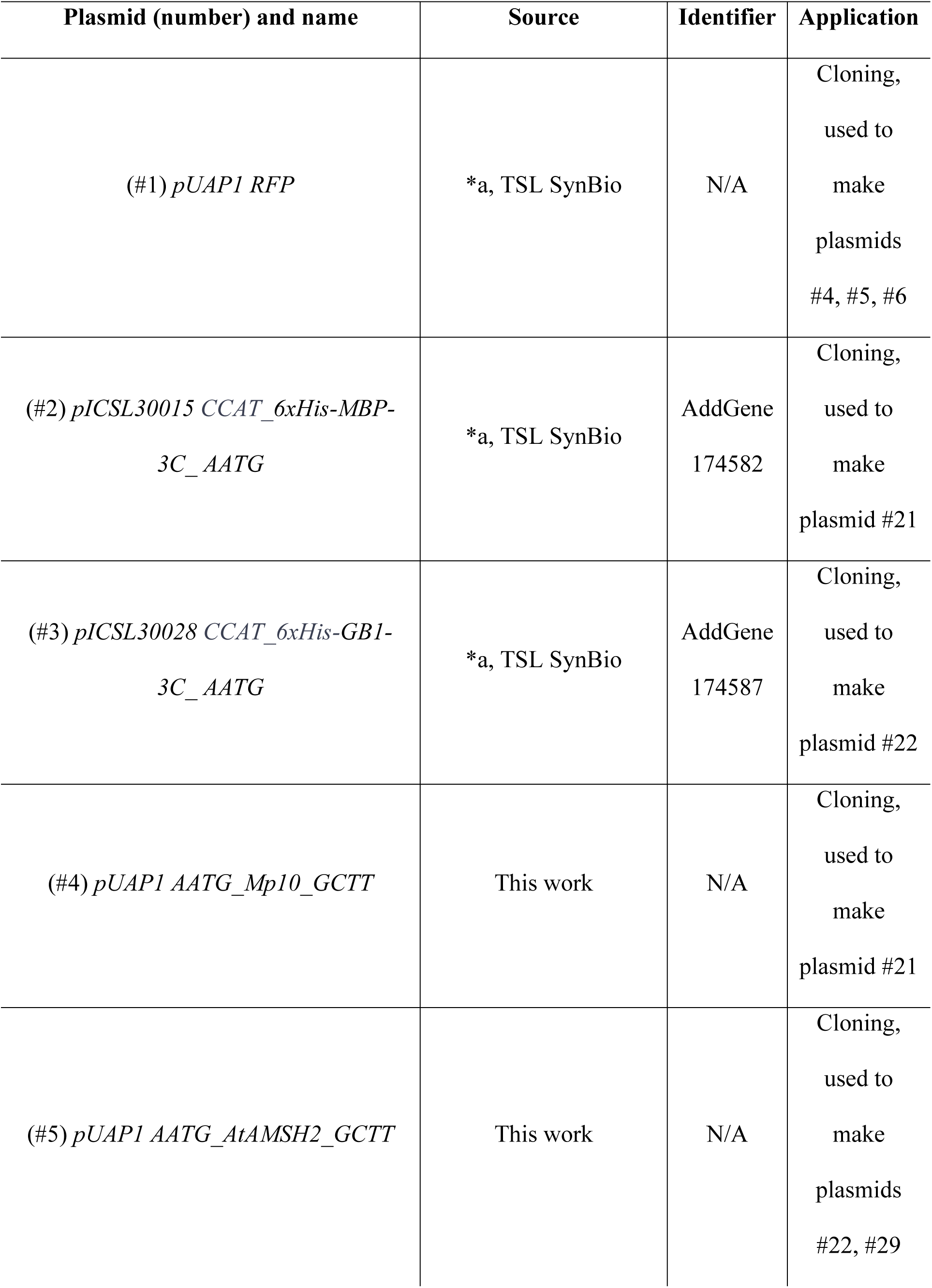
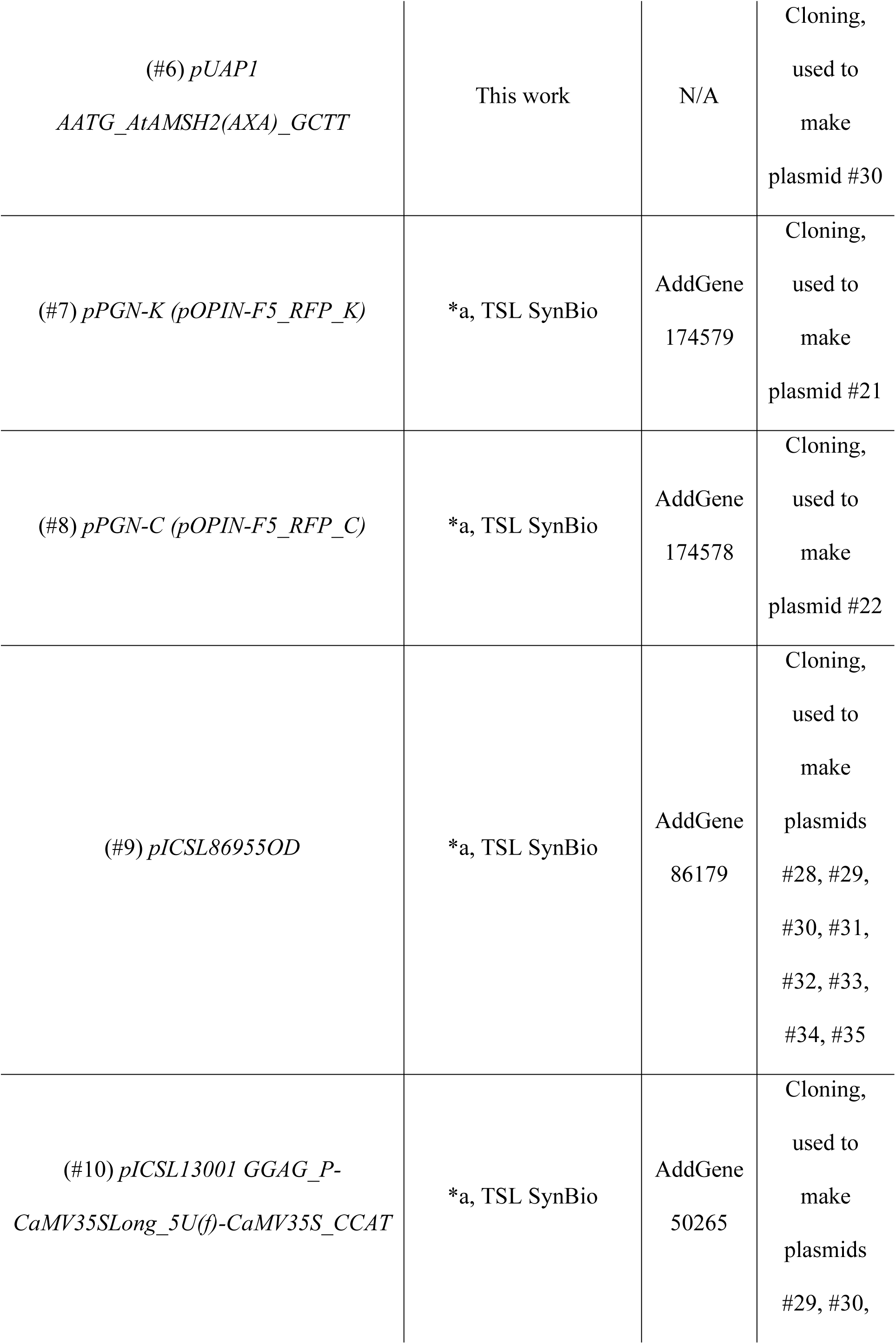
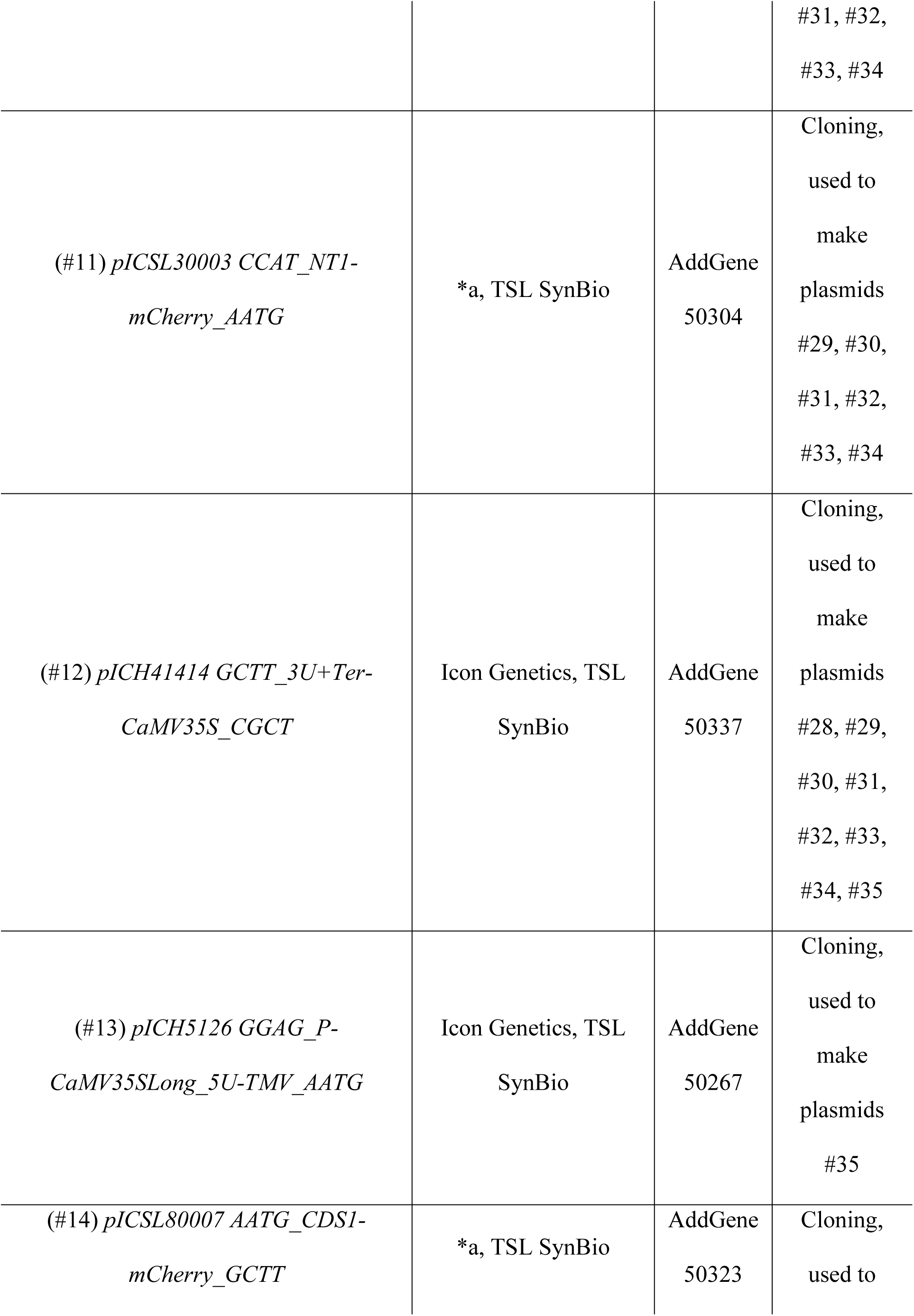
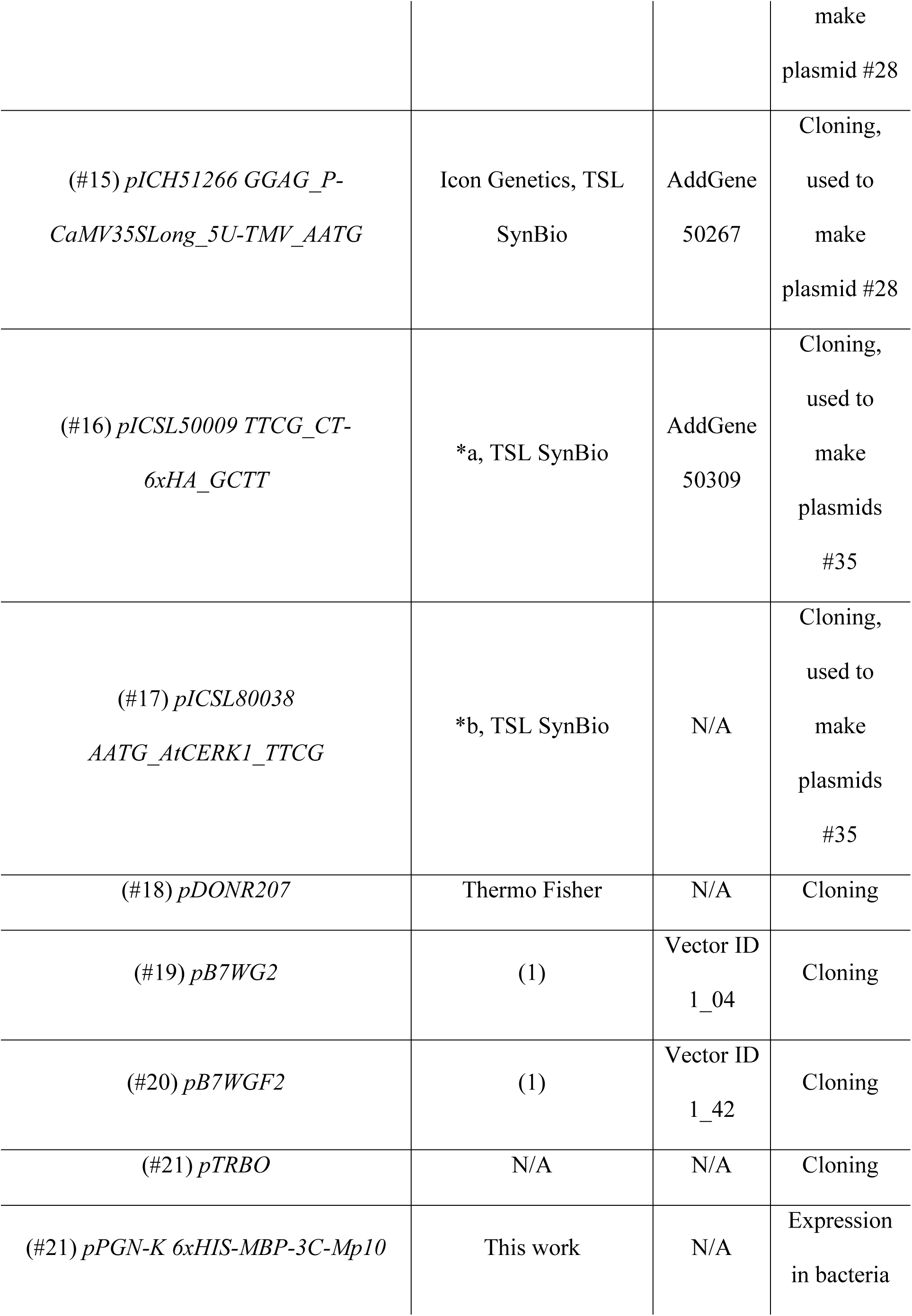
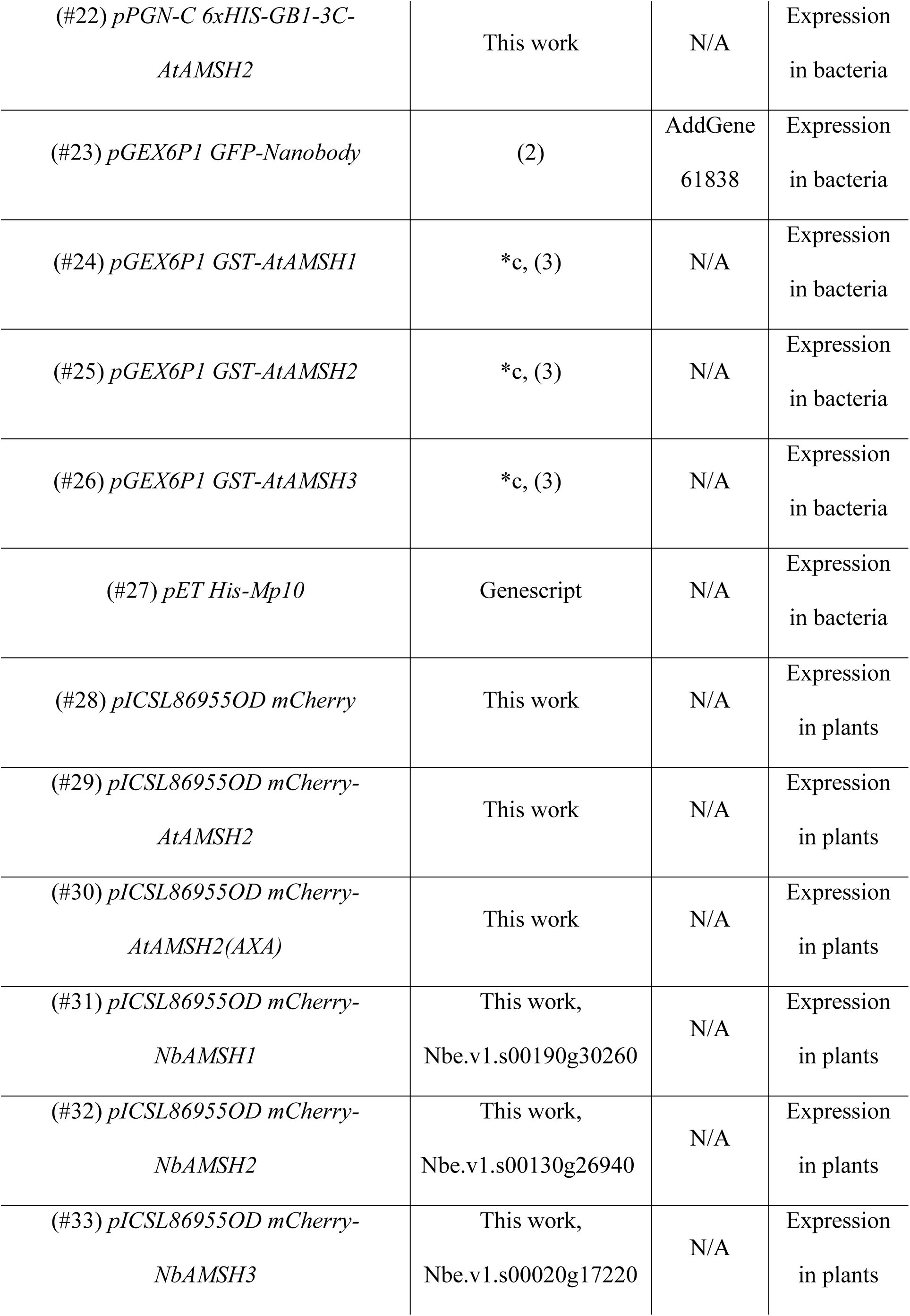
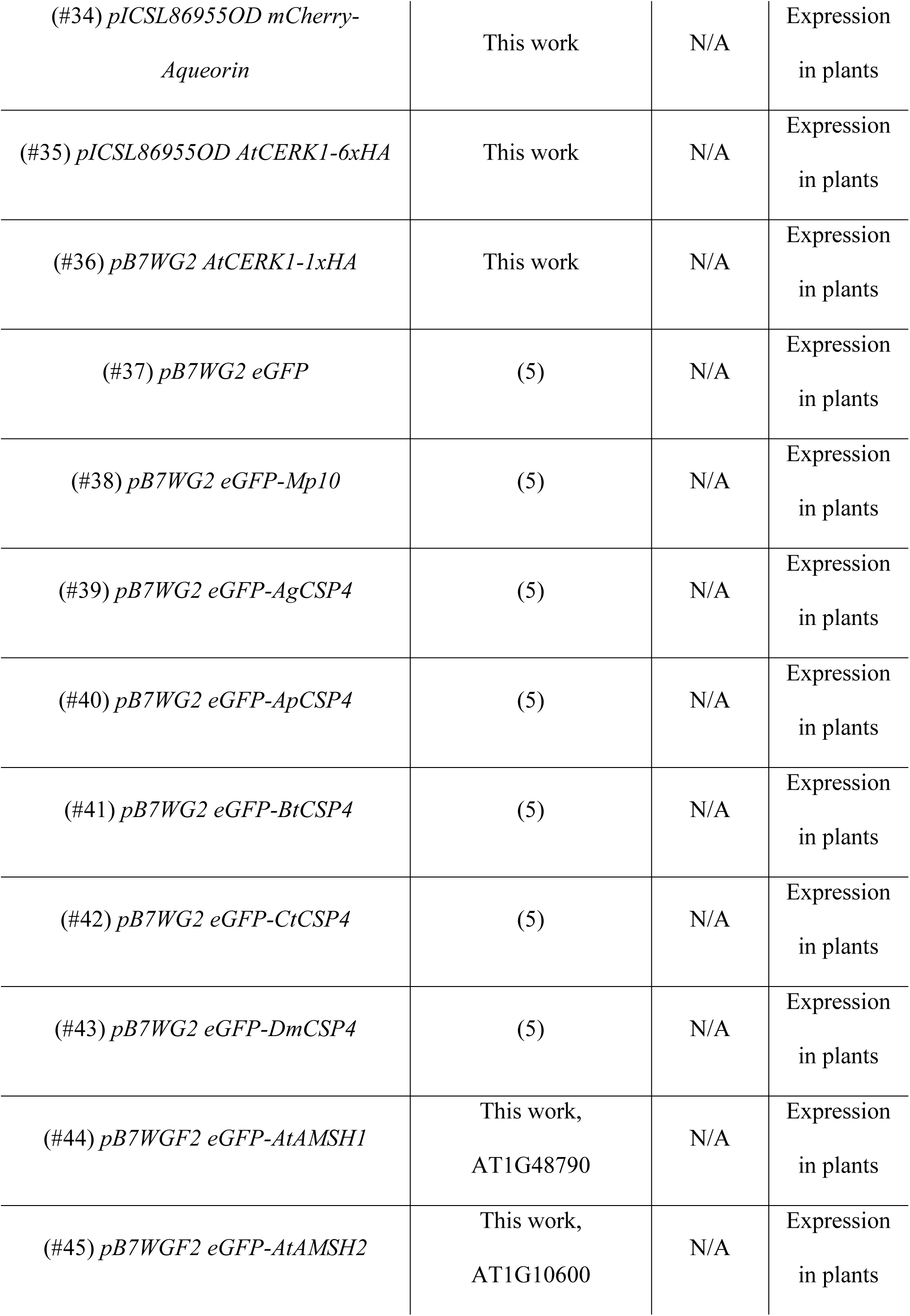
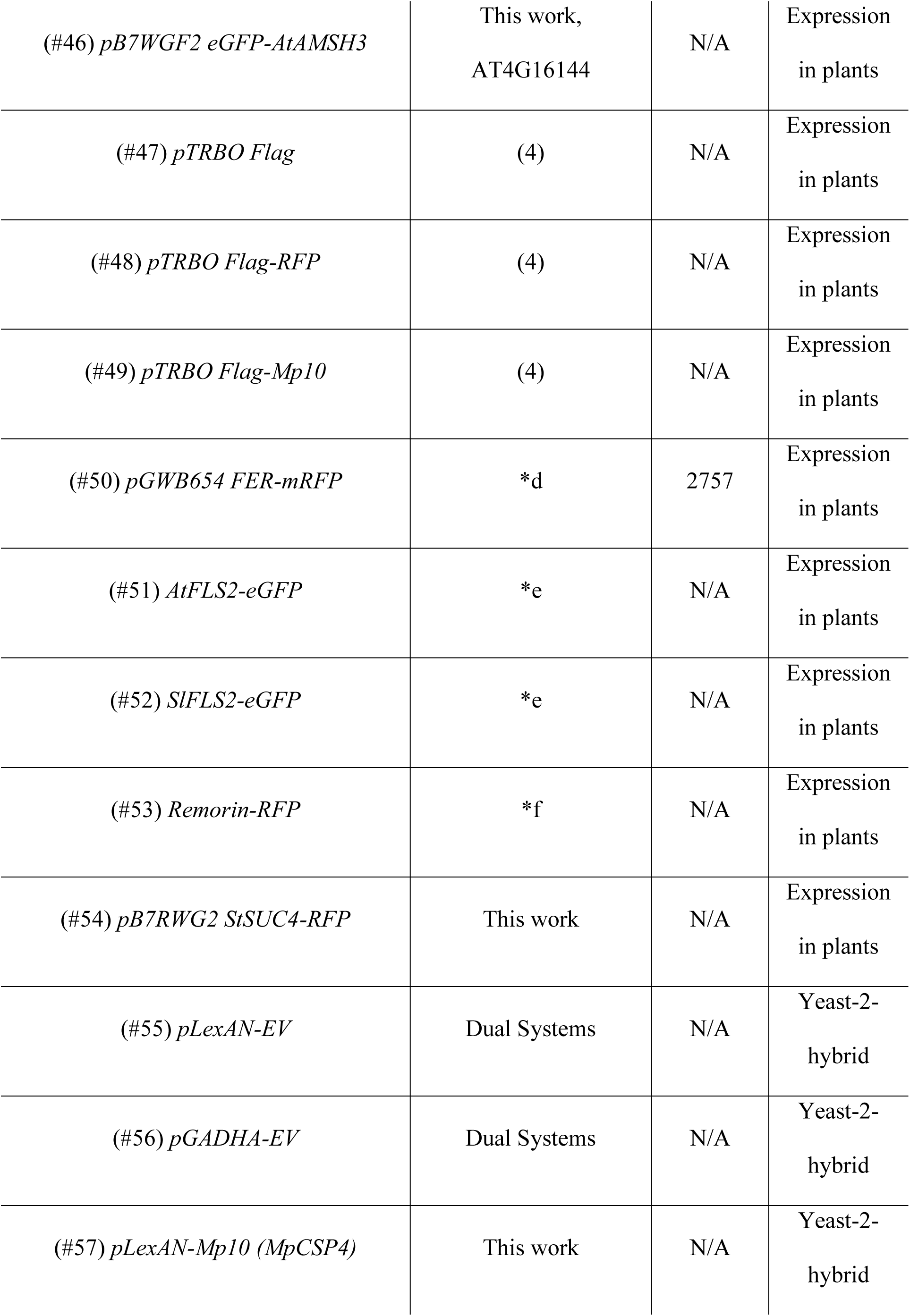
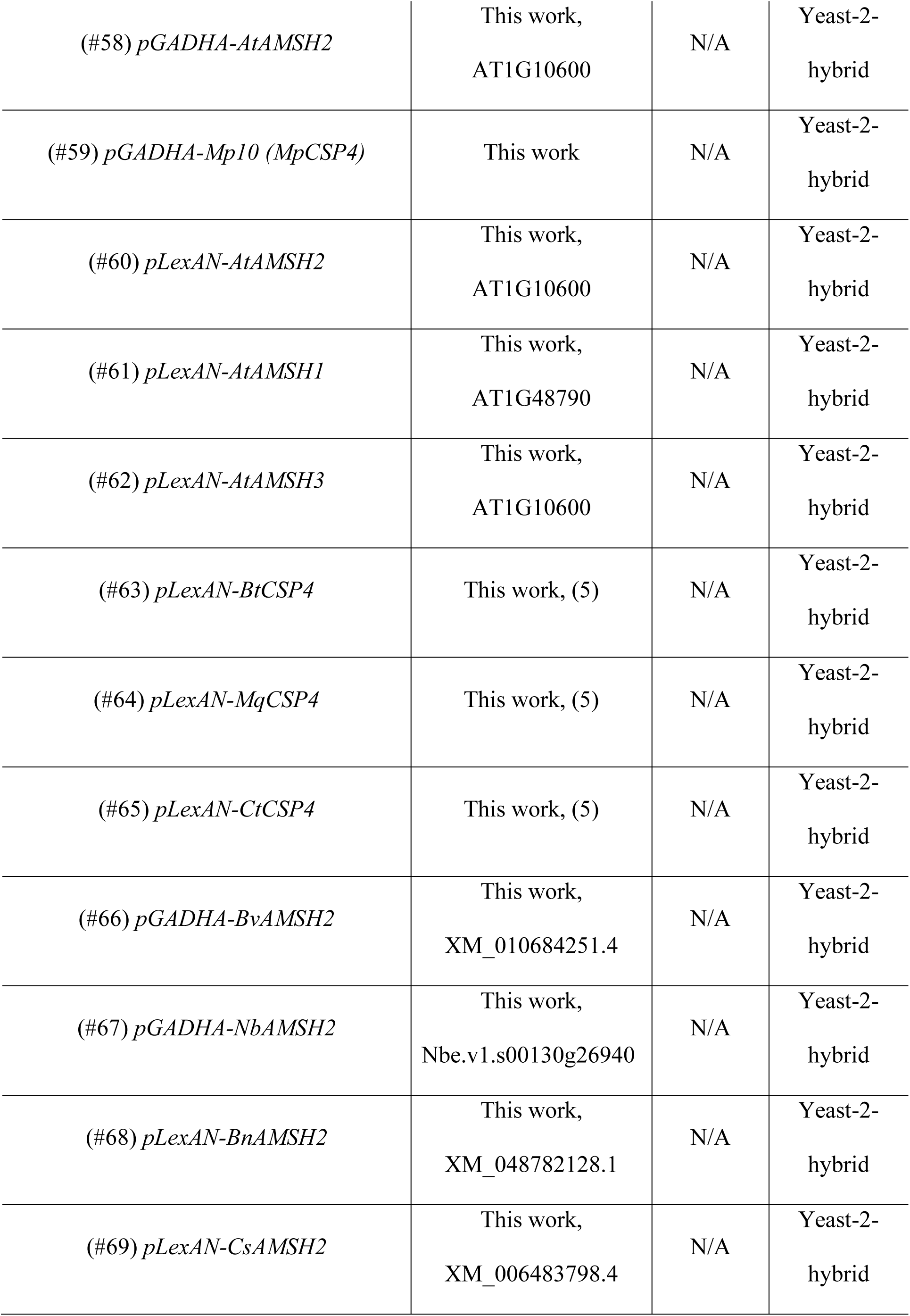
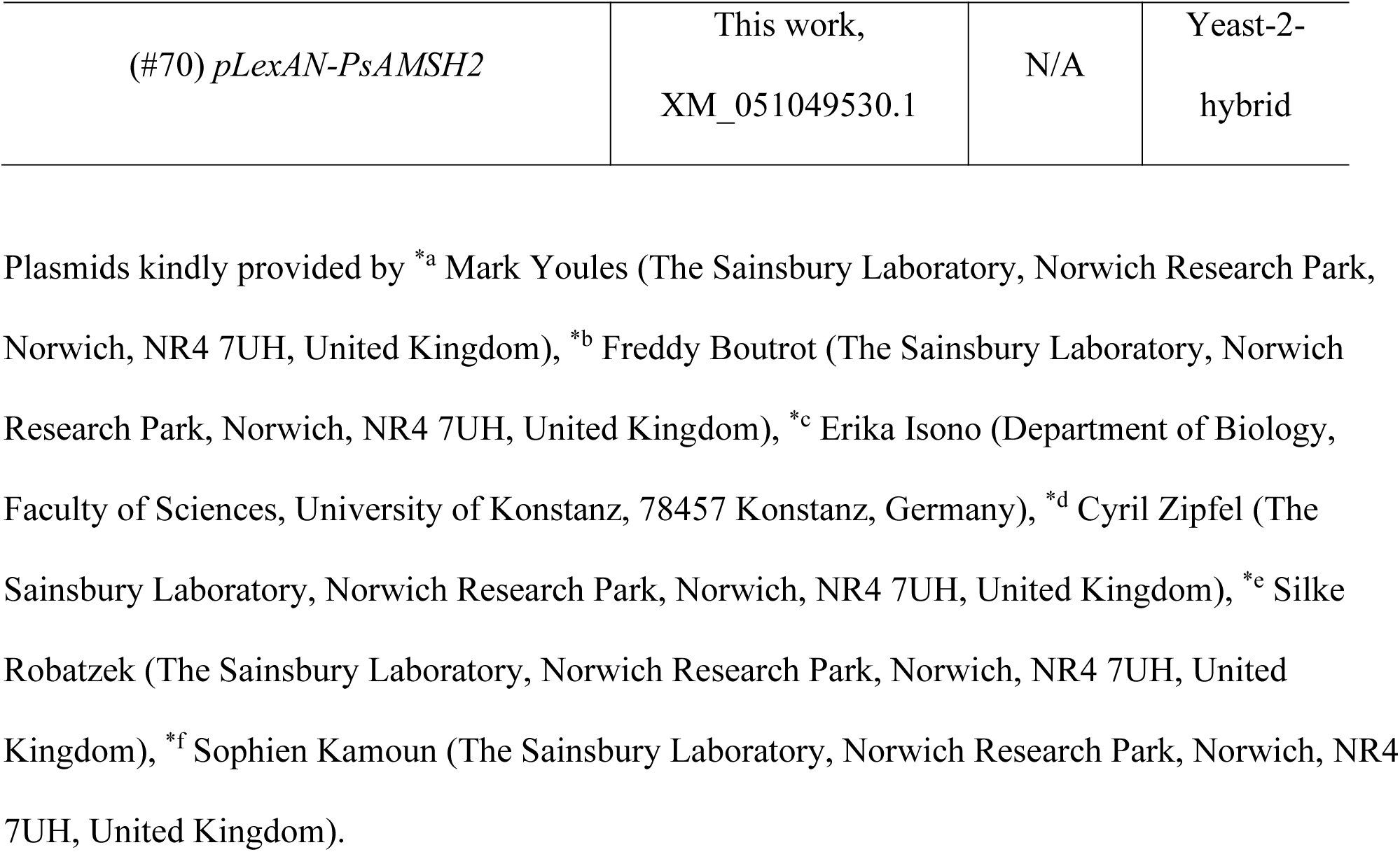
List of plasmids used in this study. The table includes annotation of all plasmids used in this study, including plasmid name, source, identifier and application.

---

**S4 Table.**
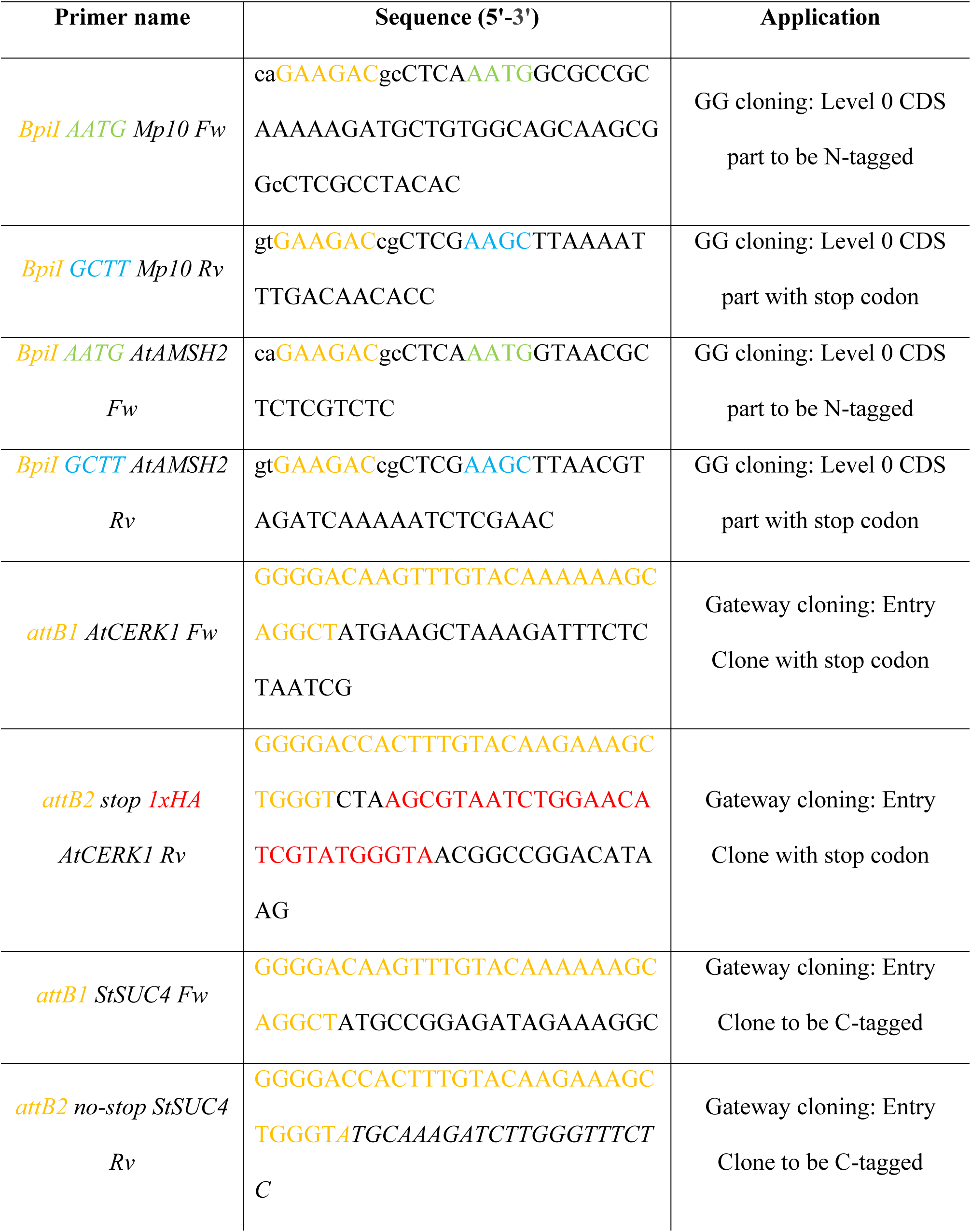
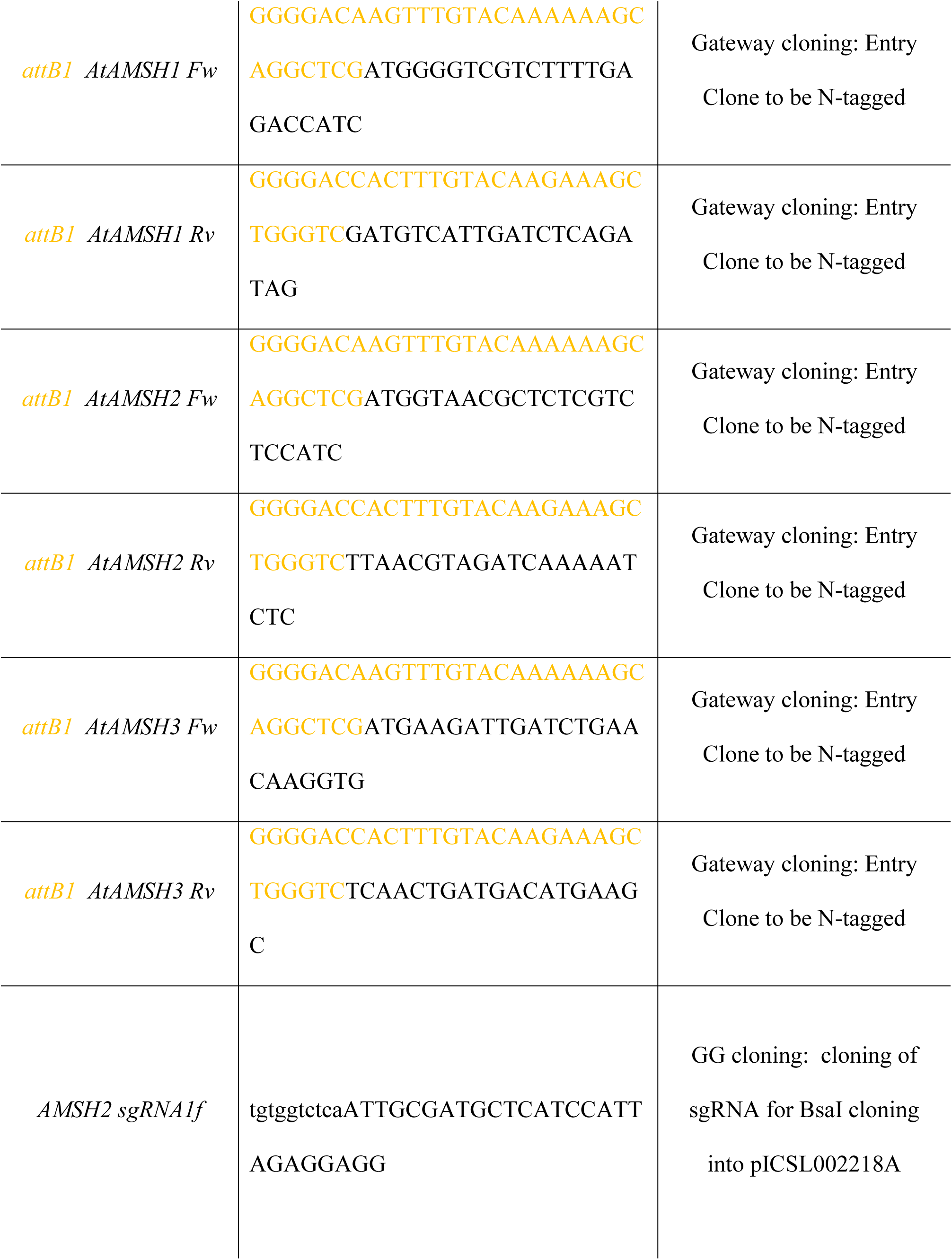
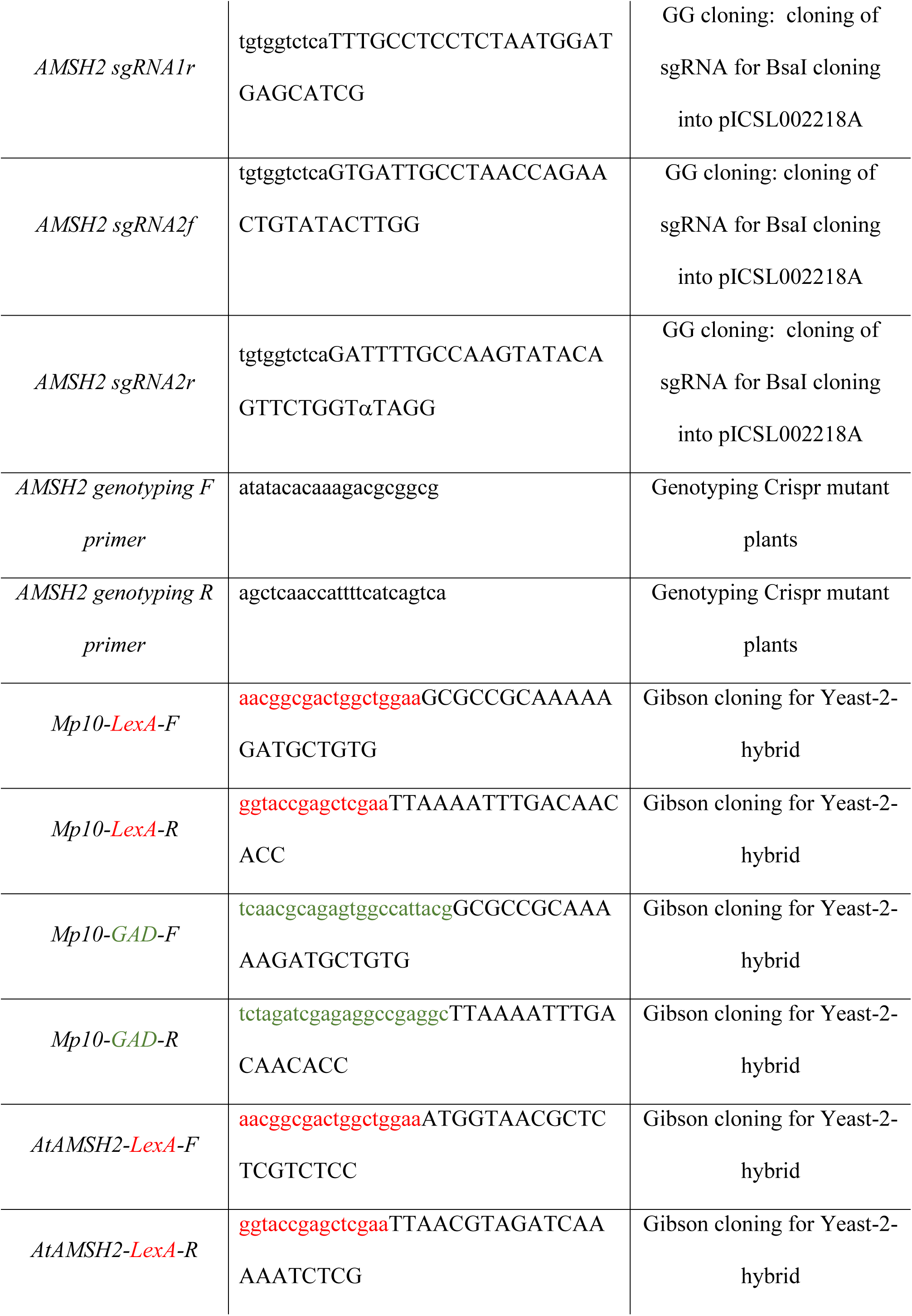
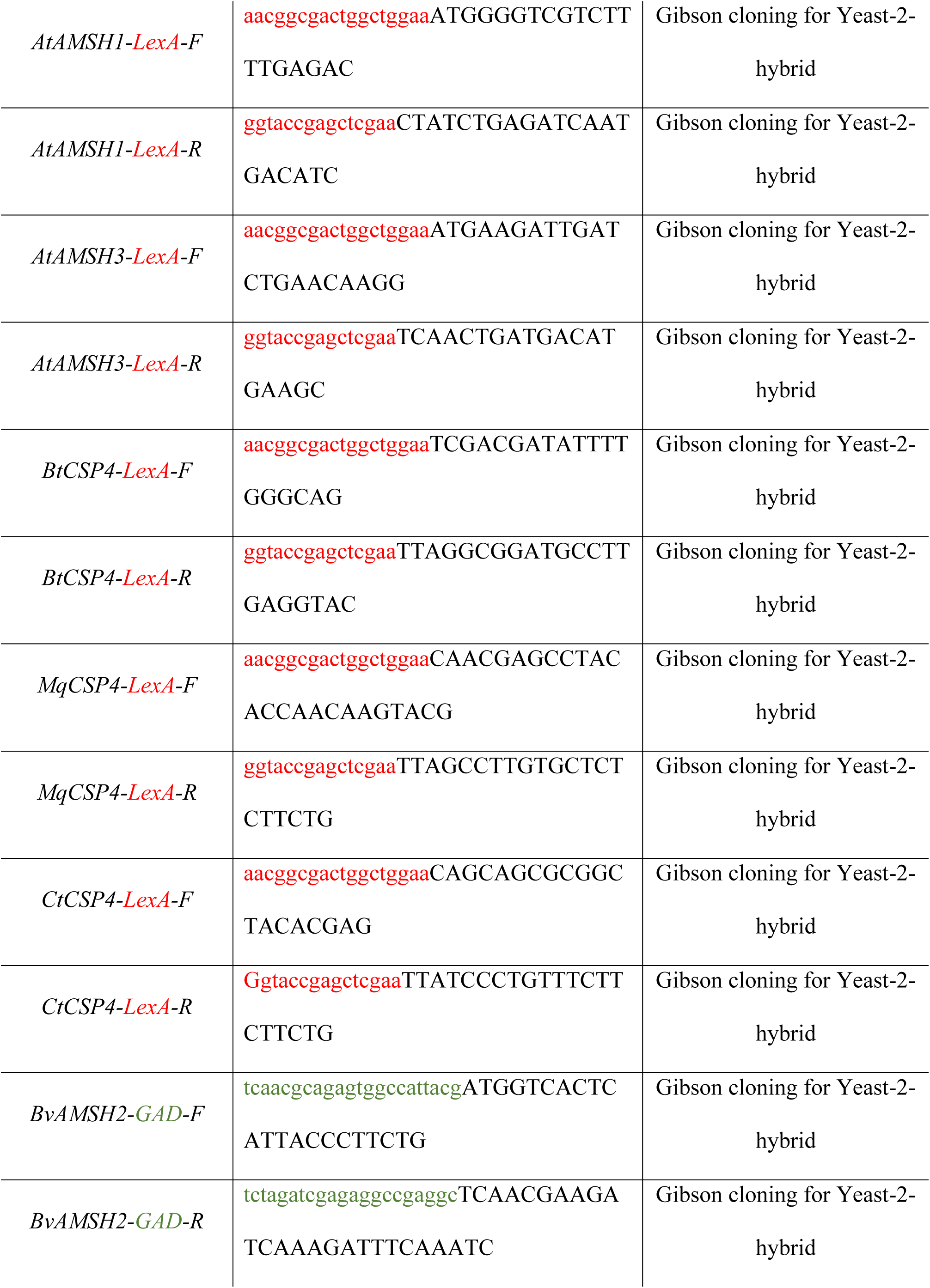
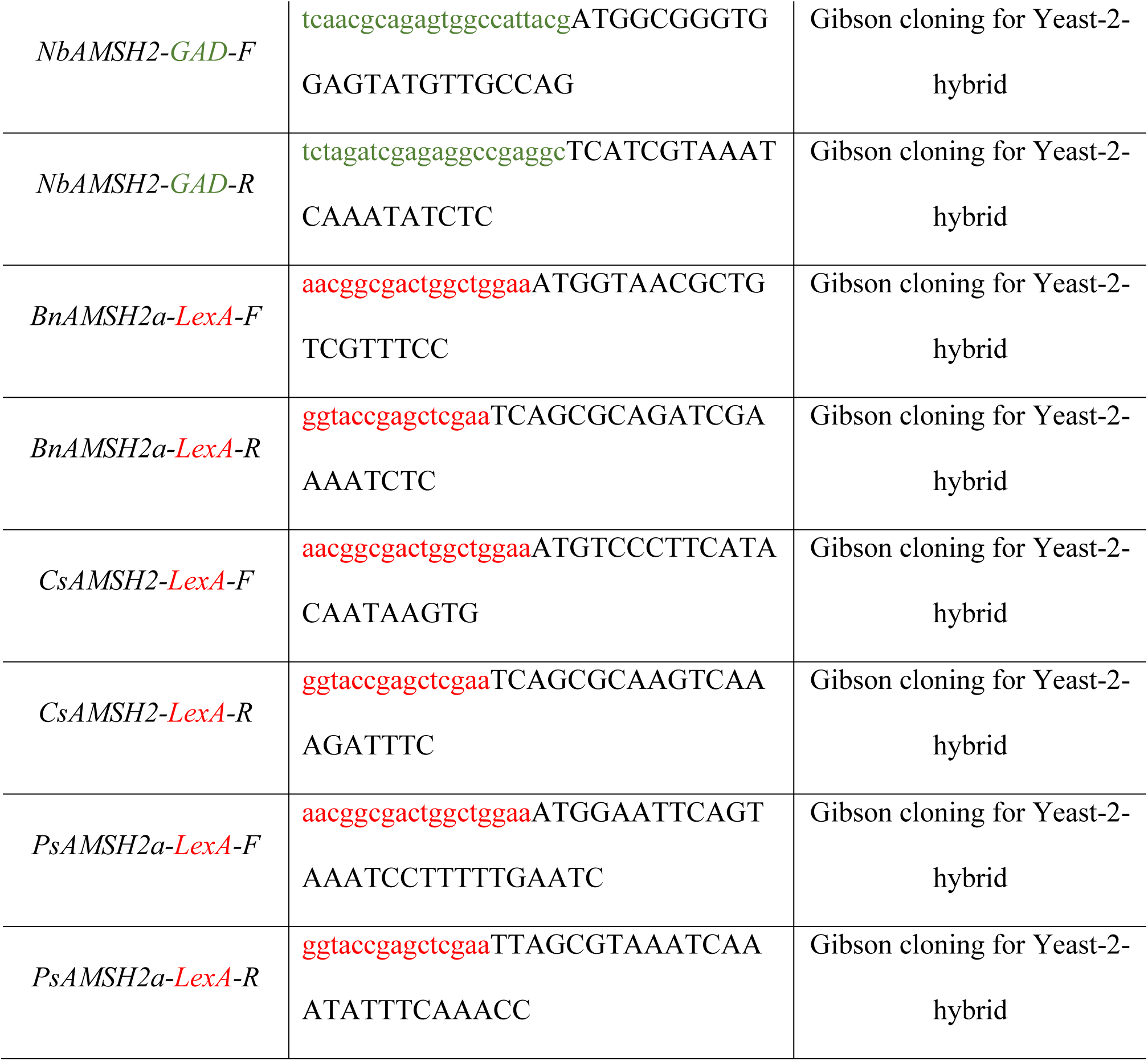
List of primer sequences used in this study. The table includes annotation of all primers used in this study, including primer name, sequence (5’-3’), and application.

**S1 Protocol. Detailed description of the LC-MS/MS methods.** The protocol includes a more detailed description of the mass spectrometry methods.

Protein pellets were resuspended in 50 µl of 1.5% sodium deoxycholate (SDC; Merck) in 0.2 M EPPS-buffer (Merck), pH 8.5 and vortexed under heating. Cysteine residues were reduced with dithiothreitol, alkylated with iodoacetamide, and the proteins digested with trypsin in the SDC buffer according to standard procedures. After the digest, the SDC was precipitated by adjusting to 0.2% trifluoroacetic acid (TFA), and the clear supernatant subjected to C18 SPE using home-made stage tips with C18 membrane plugs (Supelco Analytical - 3M, Bellafonte, PA). Aliquots were analysed by nanoLC-MS/MS on an Orbitrap Eclipse™ Tribrid™ mass spectrometer equipped with a FAIMS Pro Duo interface and coupled to an UltiMate^®^ 3000 RSLCnano LC system (Thermo Fisher Scientific, Hemel Hempstead, UK). The samples were loaded onto a trap cartridge (PepMap™ Neo Trap Cartridge, C18, 5um, 0.3×5mm, Thermo) with 0.1% TFA at 15 µl min^-1^ for 3 min. The trap column was then switched in-line with the analytical column (Aurora Frontier TS, 60 cm nanoflow UHPLC column, ID 75 µm, reversed phase C18, 1.7 µm, 120 Å; IonOpticks, Fitzroy, Australia) for separation at 55°C using the following gradient of solvents A (water, 0.1% formic acid) and B (80% acetonitrile, 0.1% formic acid) at a flow rate of 0.26 µl min^-1^ : 0-3 min 0% B (parallel to trapping); 3-10 min increase B (curve 4) to 7%; 10-100 min linear increase B to 32%, 100-148 min increase B to 50%; followed by a ramp to 99% B and re-equilibration to 0% B, for a total of 180 min runtime. For a second set of samples total run time was reduced to 140 min with a similar gradient. Mass spectrometry data were acquired with the FAIMS device set to three compensation voltages (-35V, -50V, -65V) at standard resolution for 1.0 s each with the following MS settings in positive ion mode: OT resolution 120K, profile mode, mass range m/z 300-1800, normalized AGC target 100%, max inject time 50 ms; MS2 in IT Turbo mode: quadrupole isolation window 1 Da, charge states 2-5, threshold 1e^4^, HCD CE = 30, AGC target standard, max. injection time dynamic, dynamic exclusion 1 count for 15 s with mass tolerance of ±10 ppm.

The mass spectrometry raw data were processed and quantified in Proteome Discoverer 3.1 (Thermo) using the search engine CHIMERYS (MSAID, Munich, Germany); all mentioned tools of the following workflow are nodes of the proprietary Proteome Discoverer (PD) software. The predicted *N. benthamiana* proteome from the Nbe_v1.1 genome assembly (https://nbenthamiana.jp/) (1) was used to identify the proteins and provide a functional description based on homologs. The Nbe_v1_pep.fasta database was imported into PD adding a reversed sequence database for decoy searches; in the same way a custom database with the Mp10 and AMSH construct sequences as well as a database for common contaminants (maxquant.org, 246 entries, Aug 2024) were also included. The CHIMERYS database search was performed with the inferys_3.0.0_fragmentation prediction model, a fragment tolerance of 0.3 Da, enzyme trypsin with 2 missed cleavages, variable modification oxidation (M), fixed modification carbamidomethyl (C) and FDR targets 0.01 (strict) and 0.05 (relaxed). The workflow included the Minora Feature Detector with min. trace length 7, S/N 3, PSM confidence high and the TopN peak filter with 10/100 Da. The consensus workflow in the PD software was used to evaluate the peptide identifications and to measure the abundances of the peptides based on the LC-peak intensities. For identification, an FDR of 0.01 was used as strict threshold, and 0.05 as relaxed threshold. For quantification, 3 or 4 replicates per condition were measured. In PD3.1, the following parameters were used for ratio calculation: chromatographic alignment and feature linking with a retention time tolerance of 2 min, mass tolerance of 2 ppm and S/N 3; normalisation on total peptide abundances; protein abundance-based ratio calculation using the top3 most abundant peptides; missing values imputation by low abundance resampling; hypothesis testing by t-test (background based); adjusted p-value calculation by BH-method. The results were exported to a Microsoft Excel table including data for protein abundances, ratios, p-values, number of peptides, protein coverage, the CHIMERYS identification score and other important values.

